# A high-quality genome of the mass-blooming desert plant *Cistanthe longiscapa* and its photosynthetic behavior related to drought and life history

**DOI:** 10.1101/2024.11.06.622337

**Authors:** Anri Chomentowska, Pauline Raimondeau, Lan Wei, Eleanor G.D. Jose, Sophie G. Dauerman, Virginia Z. Davis, Andrés Moreira-Muñoz, Iris E. Peralta, Erika J. Edwards

## Abstract

- Crassulacean acid metabolism (CAM) photosynthesis has independently evolved many times in arid-adapted plant lineages. *Cistanthe longiscapa* (Montiaceae), a desert mass-blooming annual, can upregulate CAM facultatively upon stress such as drought. Few studies, however, consider life history stages when measuring CAM activity or its facultative onset.
- To test the effect of drought and flowering on photosynthetic activity, we assayed *Cistanthe* individuals in fully-watered and drought conditions, as well as fully-watered individuals at pre-flowering and flowering life stages. We assembled and annotated a chromosome-scale genome of *C. longiscapa* and compared it with the genome of *Portulaca amilis* and analyzed differential gene expression.
- Results show significantly upregulated CAM in drought conditions as compared to fully-watered conditions; furthermore, flowering individuals showed slightly higher CAM activity as compared to pre-flowering plants, even when fully-watered. Differential gene expression analyses provide preliminary support for the possible co-regulation of CAM expression and reproduction.
- We emphasize the potentially missed significance of life history in the CAM literature, and consider how the CAM biochemical module could become co-opted into other plant behaviors and responses, such as the shift to reproduction or flowering in annuals.

## Introduction

Ephemeral organisms, such as desert annuals in plants, harbor adaptations unique to their life history strategy. For example, annuals may choose to allocate their resources differently upon exposure to stress, as compared to long-living organisms, given that they must reproduce before death in a short period of time (Roumet *et al*., 2006; Thomas, 2013; Lundgren & Des Marais, 2020). Annual plants are often associated with a drought-escape strategy (i.e., only emerging after rainfall and reproducing before the depletion of water supply; Ludlow, 1989), but in reality annuals can and do experience drought at various stages of life depending on the desert environment (Eshel *et al*., 2021). For certain lineages of desert plants, a drought adaptation that they may utilize in resource maximization is Crassulacean Acid Metabolism (CAM) photosynthesis.

CAM photosynthesis is characterized by the initial nocturnal fixation of CO_2_ in the form of a four-carbon acid and its daytime decarboxylation for the final fixation of CO_2_ by RuBisCO. This temporal separation of carbon intake and final fixation allows CAM plants to minimize transpirational water loss, since the stomata are open at night when temperatures are cooler and thus, evapotranspiration rates are lower (Neales *et al*., 1968; Osmond, 1978), thus making CAM plants the most water-use efficient plants on Earth (Cockburn, 1985; Nobel, 1996). A major enzyme in this pathway is phosphoenolpyruvate carboxylase (PEPC), which catalyzes the fixation of atmospheric CO_2_ into oxaloacetate and then eventually malate, which is stored in the vacuole overnight as malic acid (Holtum *et al*., 2005). During the day, stomata remain closed, and the decarboxylation of malate creates a high CO2-concentration in photosynthesizing tissue, allowing Rubisco efficient final carboxylation via the Calvin-Bensen-Bassham cycle (Lüttge, 2002, 2011). The fixation of carbon using the CAM cycle does come with energetic costs, with models predicting three times higher nocturnal requirements of ATP and reducing agents in CAM compared to C3, though the costs are mostly alleviated by lower photorespiration rates in CAM plants due to their elevated levels of internal daytime CO2 concentrations (Shameer *et al*., 2018). CAM photosynthesis has independently evolved many times in diverse lineages across the plant phylogeny, with estimates of up to 7% of all vascular plants having a functional CAM cycle (Gilman *et al*., 2023).

An important aspect of CAM biology is the diversity of ways in which the biochemical cycle has been integrated into a plant’s physiology. In many canonical desert succulent lineages such as cacti, aloes, and euphorbias, CAM has become the plant’s primary photosynthetic metabolism, operating daily and under varied conditions (termed “strong CAM” in Edwards 2019, “primary CAM, or pCAM” in Gilman *et al*. 2024). There are likely even more species, however, that use CAM as an accessory metabolism and still fix the majority of atmospheric CO_2_ directly by Rubisco. Edwards (2019) provided an umbrella term “C3+CAM” to refer to all of these phenotypes (“minority CAM, or mCAM” of Gilman *et al*., 2024), but the phenotypes are quite diverse. They include a weakly expressed constitutive CAM, a fully inducible and reversible facultative CAM, an upregulated CAM cycle that is apparently not reversible, and CAM-cycling and CAM-idling phenotypes, which use the pathway only to refix internal respiratory CO_2_ (Winter, 2019). All of these behaviors can be found in both annual and perennial plants with one notable exception: there are no known annual plants that are also strong CAM species.

*Cistanthe longiscapa* (Phil.) Peralta & D.I. Ford is an annual C3+CAM species in the Montiaceae (Portulacineae; Caryophyllales; Fig. 1c), native to the Atacama desert in northern Chile (Fig.1b). *C. longiscapa* is of particular interest as a major element of the massive “superblooms’’ (*desierto florido*) that occur in the Atacama Desert, when many thousands of dormant seeds germinate simultaneously—an occurrence usually associated with El Niño years (Vidiella *et al*., 1999; Jaksic, 2001; Chávez *et al*., 2019), flowering synchronously to paint the desert floor magenta (Fig.1a). Holtum *et al*. (2021) found that *C. longiscapa* performs low-level constitutive CAM, with the ability to upregulate CAM upon drought. Importantly, for desert annuals like *Cistanthe longiscapa* that experience little to no rainfall throughout their development (Eshel *et al*., 2021; Ossa *et al*., 2022), drought stress should be most prominent in later stages of life. It is quite likely that, in the field, *C. longiscapa* is experiencing significant water deficits while it is flowering, therefore, upregulated CAM expression and flowering/fruiting should be temporally correlated during its life history. At the same time, the expression of a CAM cycle during flowering could potentially benefit an annual plant independently of its water-use efficiency, as increased carbon fixation during this time will maximize its resources to allocate to developing seeds. Maturing quickly, expending large reproductive efforts, and producing many offspring in unstable environments are characteristics of *r*- selected species in life history theory (Stearns, 1976), which applies to desert annuals such as *C. longiscapa*.

**Figure 1.**
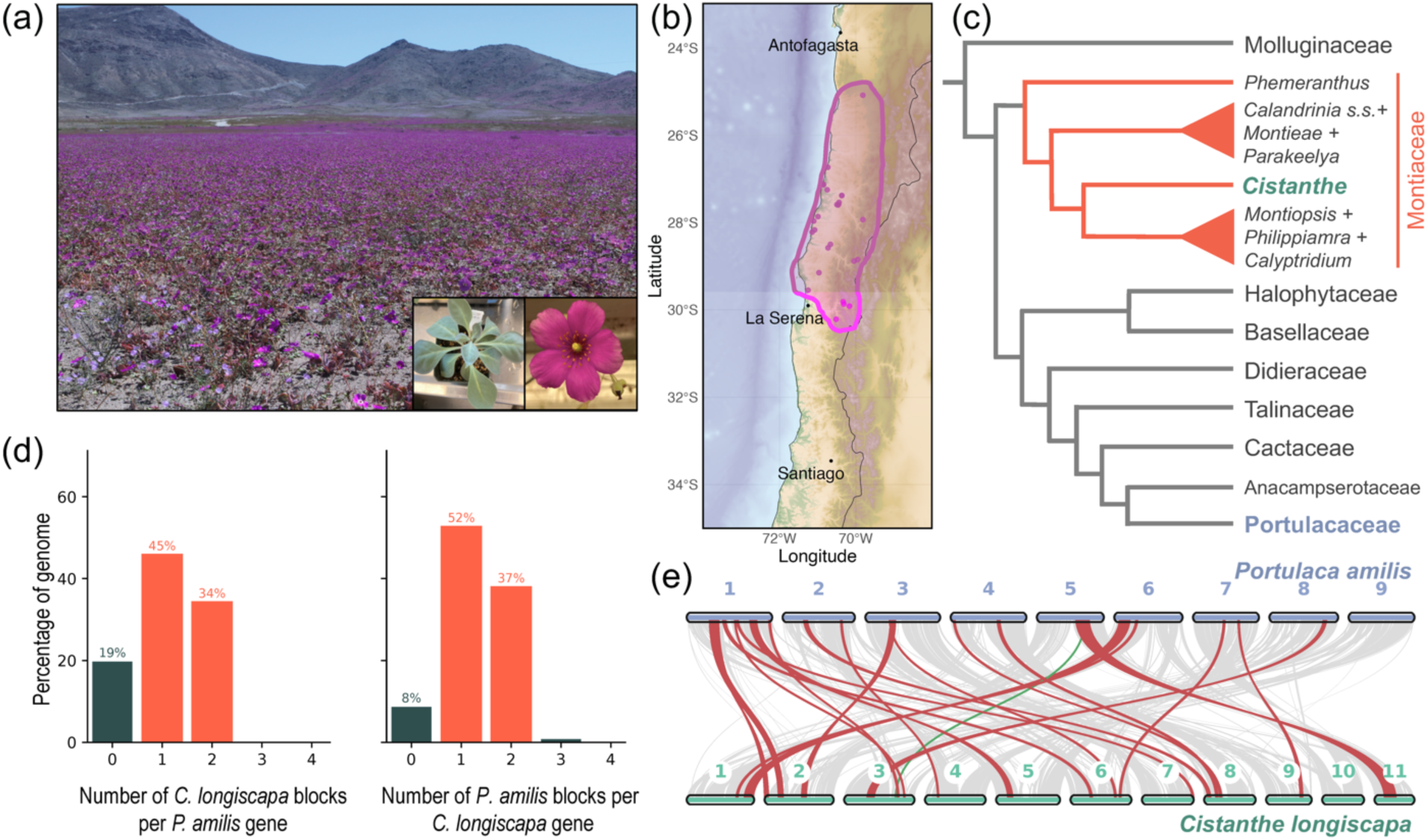
Overview of *Cistanthe longiscapa* and its genome. (a) *C. longiscapa* is a desert annual C3+CAM plant that exhibits a mass-blooming behavior in years with higher-than-usual precipitation, (b) native to the Atacama Desert in Northern Chile. Points on the map are filtered occurrence data from the Global Biodiversity Information Facility (GBIF, 2022). (c) The genus *Cistanthe* is in the family Montiaceae, highlighted in orange branches of a cladogram of the suborder Portulacineae and sister lineage Molluginaceae. (d) Syntenic depth analyses of *C. longiscap*a and *Portulaca amilis* genomes shows that 34% and 37% of the *P. amilis* and *C. longiscapa* genome had two syntenic regions, respectively. This suggests that these taxa share an ancestral whole genome duplication. (e) The syntenic blocks between the two genomes that include core CAM genes are visualized in red, whereas the rest are visualized in gray. One gene that did not map to a similar region to the rest of its block from *P. amilis* was PPC-1E1c (visualized in green), the copy of the phosphoenolpyruvate carboxylase (PEPC) gene used in CAM in *C. longsicapa* and *P. amilis*.

In plants with a facultative CAM cycle, most experiments focus on environmental factors such as drought, salinity, and occasionally photoperiod as the variables that trigger an upregulation of CAM expression (reviewed in Winter & Holtum, 2014). In contrast, very little attention has been paid to endogenous signaling or the incorporation of CAM into particular plant developmental stages or behaviors (but see Cushman *et al*., 1990; Herppich *et al*., 1992; Winter & Holtum, 2007; Bangal & Lambrecht, 2010; Ramírez & Herrera, 2017; Winter, 2019). In the only published experiment of *Cistanthe longiscapa*, drought induced an upregulation of CAM activity, but re-watering did not reverse the CAM behavior—it maintained an elevated CAM expression throughout the experiment, which included over 30 days of continuous gas exchange monitoring (Holtum *et al*., 2021). Though not reported, the plant was likely flowering and fruiting during this time, suggesting a possible connection between continued CAM expression and a shift to reproduction.

In this study we investigate a potential additional phenotype to include in the “C3+CAM” umbrella, which we term here “reproductive CAM”. In annual plants with a functional CAM cycle, it could be advantageous to express that cycle even in the absence of drought, provided the plant is receiving sufficient light to fuel the energetic requirements of the CAM cycle. Here, we present *Cistanthe longiscapa* as an excellent new experimental system to investigate reproductive CAM and other C3+CAM behavior. We sequenced and annotated a chromosome-level genome, characterized its growth trajectory, and performed a series of experiments designed to induce CAM under drought and also to monitor CAM under well-watered conditions during the transition to flowering and fruiting. We demonstrated increased expression of CAM in reproductive phases under well-watered conditions, indicating some degree of reproductive CAM. Gene expression analyses mostly support our physiological measurements, and motif-enrichment analyses demonstrate shared transcription factor binding sites with known CAM genes and other gene clusters upregulated during flowering/fruiting, suggesting that the CAM pathway may be co-regulated with reproduction. We discuss different hypotheses for how reproductive CAM could evolve from a drought-induced facultative CAM phenotype.

## Materials and Methods

### Plant materials and growth conditions

*Cistanthe longiscapa* specimens were collected ∼20 km south of Copiapó, Atacama region, Chile (27.521583 S, 70.432833 W, elevation 730 m) and propagated at James Cook University, Australia. Seeds from the propagation were cold-stratified then germinated on regular potting soil at Yale University. Half were cultivated in a 12-hour photoperiod chamber, the other half in a 14-hour photoperiod chamber. Plants in both chambers were exposed to 200 µmol of light during the photoperiods, and grown under ambient CO2. The temperature was maintained at 25 ℃ during the day and 20 ℃ at night. The relative humidity was maintained such that it was the same inside and outside the growth chambers.

Leaf tissue from the juvenile stage (after a 72-hour dark treatment) was collected and flash frozen with liquid nitrogen, then sent to the Genomics and Cell Characterization Core Facility (GC3F) at University of Oregon for high molecular weight (HMW) DNA extraction and subsequent PacBio HiFi library preparation and sequencing. Leftover material was used for Hi-C library preparation and sequencing. Furthermore, six samples of leaf, root, bud, and flower tissue (approx. 0.3 ∼ 0.5 g total) were collected both in the day and nighttime and flash frozen. Total RNA was extracted and Iso-Seq libraries were prepared from these samples to sequence full-length mature mRNA from the whole transcriptome.

### Life history and gas exchange measurements

To characterize the typical lifespan and growth of *Cistanthe longiscapa*, we examined a total of 115 plants split between the 12- and 14 hour chambers every two days to count the number of leaves, buds, flowers, and fruits. The dates of germination, reproductive stalk emergence, and flowering were recorded. In the 14-hour chamber, we also measured gas exchange in one pre-flowering plant under a well-watered condition and one under drought, to check that the plant was in fact showing a physiological response to our drought condition. We used the LI-6800 Portable Photosynthesis System (LI-COR Biosciences) with a clear-top chamber, and spot-measured gas exchange in leaves across 6 timepoints (4-hour intervals). The CO_2_ assimilation rate (µmol m^-2^ s^-1^) was adjusted for the total leaf area in the chamber head.

### Genome and whole-transcriptome sequencing

To assemble a high-quality genome, we chose an approach that allowed for sequencing long-reads and scaffolding at the chromosome level: PacBio HiFi and Hi-C sequencing. HMW DNA extraction and library preparation was performed at University of Oregon (GC3F). The genomic library was sequenced across three PacBio Sequel II SMRT cells based on prior genome-size estimates from flow cytometry (1.35 Gbp +/- 10%). The Phase Genomics Hi-C library was also prepared at GC3F, and sequenced on the Illumina NovaSeq 6000 (2 × 150-bp paired-end reads).

Additionally, to obtain full-length transcripts for high-quality annotation of the genome, we sequenced the whole-transcriptome with Iso-Seq. For this, we extracted high-quality RNA from six samples of flash frozen tissue by grinding them with mortar and pestle in liquid nitrogen, and using a modified CTAB/ TRIzol/sarkosyl-based protocol from Jordon-Thaden *et al*. (2015; ‘Option 2’). This resulted in an average yield of 34000 ng of total RNA across samples. The Yale Center for Genome Analysis (YCGA) prepared an Iso-Seq library from these extractions, and sequenced the library on a PacBio Sequel II flow cell.

### Genome assembly and annotation

Subreads from HiFi sequencing were collapsed into Circular Consensus Sequence (CCS) reads before demultiplexing/quality-filtering with the PacBio *CCS* 6.4.0 (https://github.com/PacificBiosciences/ccs) pipeline. We used HiFiasm v0.19.8 (Cheng *et al*., 2021, 2022) to generate a de-novo assembly combining HiFi and Hi-C reads, and YAHS (Zhou *et al*., 2023) for further scaffolding. Hi-C reads were mapped to the draft assembly using BWA-MEM (parameters -A1 -B4 -E50 -L0; Li, 2013). The scripts ‘perl filter_five_end.pl’ and ‘perl two_read_bam_combiner.pl’ from Arima Genomics Mapping Pipeline v03, as well as SAMtools (Danecek *et al*., 2021), were used to process and sort the Hi-C read alignments. We used Juicer (Durand *et al*., 2016) to generate Hi-C contact maps, scripts from https://bitbucket.org/bredeson/artisanal/src/master/src/ to prepare Hi-C map visualization, and Juicebox (Robinson *et al*., 2018) to further edit the scaffolds. Assembly completeness was assessed with BUSCO v5.7.0 against eudicots_odb10.1 (Manni *et al*., 2021).

Iso-Seq raw reads were pre-processed with the Pacbio Iso-Seq pipeline to generate CCS reads and filtered for full length non-chimeric (FLNC) reads. We converted the FLNC reads into FASTQ format with SAMtools (Danecek *et al*., 2021) and mapped them with Minimap2 using the Iso-Seq preset (Li, 2018). The final assembly was soft-masked for repeats using Red (Girgis, 2015) and annotated using Braker3 (Gabriel *et al*., 2024) with hits from OrthoDB v10 for Viridiplantae (Kriventseva *et al*., 2019) and the mapped Iso-Seq data. Functional annotation was performed with InterProScan v5.67-99 (Jones *et al*., 2014). We used EDTA (Ou *et al*., 2019) to identify transposable elements (TEs).

### Comparative genomic analyses

We compared the annotated genome of *Cistanthe longiscapa* to the most closely related published genome, C4+CAM species *Portulaca amilis* (Gilman *et al*., 2022), to assess the degree of synteny between the two species at least ∼45 million years divergent (Arakaki *et al*., 2011) using MCscan (Wang *et al*., 2012) and the jcvi python library (Tang *et al*., 2024). We used Orthofinder (Emms & Kelly, 2019) to perform orthology inference and identify orthologs for all photosynthetic genes. We collected proteomes for 11 angiosperm species (*P. amilis*, *Ananas comosus*, *Amaranthus hypochondriacus*, *Arabidopsis thaliana*, *Brachypodium distachyon*, *Helianthus annuus*, *Kalanchoë fedtschenkoi*, *Phalaenopsis equestris*, *Sorghum bicolor*, *Vitis vinifera*, *Zea mays*) and clustered them with *C. longiscapa* protein sequence using DIAMOND (ultra-sensitive mode, option ‘*MSA*’; Buchfink *et al*., 2021). We checked that the resulting species tree was congruent with the currently accepted phylogenetic relationships and used the defined orthogroups to identify homologs in *C. longiscapa*. We also inspected the trees manually to assess orthology between specific copies more precisely. In addition, we retrieved coding sequences from a denser sampling of CAM and non-CAM lineages for numerous available photosynthetic genes from Gilman *et al*. (2022). We used blastn (BLAST+ suite; Camacho *et al*., 2009) to identify homologs for these genes in *C. longiscapa* and aligned them with MAFFT version 7 (Katoh & Standley, 2013). Gene family trees were inferred using IQtree version 2 (Minh *et al*., 2020).

### CAM experimental setup and collections

We set up two experiments: 1) Comparing CAM expression levels in pre-flowering plants (vegetative or with reproductive stalk but no open flowers) between drought and well-watered conditions, and 2) comparing CAM expression levels between pre-flowering and post-flowering stages (at least one flower is open, fruiting or not) under well-watered conditions. After germination, *Cistanthe longiscapa* grows vegetatively, putting out leaves in its rosette; once a relatively consistent number of leaves accumulate, the plant will produce an indeterminate reproductive stalk (Fig. S2). Once flower buds develop and open, the plant will spend a significant portion of its life putting out more flowers and developing many fruits simultaneously until death. Therefore, we binned its life history between two stages, pre-flowering and post-flowering, representing a clear difference in the plant’s life history stages. The first experiment was conducted in the 12-hour photoperiod chamber, and the second was conducted in the 14-hour chamber. To induce a drought response, plants were withheld water for 10 ∼ 14 days. There were at least four biological replicates for each of the treatment x life history stage x photoperiod categories from the two experiments. Leaf tissue was flash frozen from all the biological replicates from six time points every four hours (14:00, 18:00, 22:00, 02:00, 06:00, 10:00) for bulk RNA extraction. Additional leaf tissue was also collected at dusk and dawn (18:00 / 06:00) for titratable acidity assays.

### Titratable acidity assays

CAM activity was quantified by assaying titratable leaf acidity at dusk and dawn and calculating its change; since malic acid is stored in the vacuole overnight and decarboxylated during the day, the difference in titratable acidity will be measurable in plants using CAM photosynthesis. To do this, we weighed and boiled 0.3 – 0.4 g of fresh leaf tissue in 60 mL of 20% EtOH until the volume was reduced to 30 mL. Volume was restored to 60 mL with distilled H2O, and the boiling and refilling process was repeated; the resulting supernatant was left to cool to room temperature. Once cooled, we titrated each sample with 0.002M NaOH to a pH of 6.5, and then to 7.0. We recorded the initial pH as well as the total microliters of the titrant (NaOH) used. Titratable acidity (in μ equivalents of H^+^) was calculated by mL volume titrant (up to pH 7.0) x Molarity (mM) of titrant, and divided by initial leaf sample weight (g), to give µmol of H^+^ per gram of fresh weight (µmol H^+^ g^-1^ FW). The difference between dawn and dusk leaf collection per sample, ΔH^+^, or the change in titratable acidity, is the measure for CAM activity.

### RNA extractions and sequencing

We extracted total RNA from 96 *Cistanthe longiscapa* leaf samples (four biological replicates for each of the six timepoints across the four treatments/life stages) as we did for IsoSeq. The leaf tissues were disrupted while frozen using TissueLyser III (Qiagen) with 3 mm stainless steel beads. Quantity and quality of the extractions were measured using NanoDrop spectrophotometer (Invitrogen by Thermo Fisher Scientific, Carlsbad, CA, USA) and Agilent Bioanalyzer v.2100 (Santa Clara, CA, USA). Six samples failed to meet the concentration and RIN score requirements (≥ RIN 8) for library preparation. The RNA libraries were prepared using Poly-A selection; here, three additional samples did not meet the required concentration post-PCR. A total of 87 libraries were sequenced on the Illumina NovaSeq X PLUS (2 × 150-bp paired-end reads) at the YCGA, requesting ∼25M reads per sample. All but one library were successfully sequenced, resulting in RNA-Seq data from 86 samples with each timepoint x treatment represented by three or four biological replicates (Table S1 for details).

### RNA-Seq assembly and abundance quantification

For all of the sequenced RNASeq samples, we used Trimmomatic v0.39 (Bolger *et al*., 2014) to trim out the adapter sequences, and FastQC (Andrews, 2010) to quality control the fragments. Then, we used STAR v2.7.11 (Dobin *et al*., 2013) to map the reads from each sample to our assembled genome (options ‘*-- outFilterMismatchNmax 2*’, ‘*--outSAMtype BAM SortedByCoordinate*’). To quantify transcript abundance per gene, we quantified the number of mapped reads to each gene using the program FeatureCounts (implemented in Subread v2.0.3; Liao *et al*., 2013, 2014) with the ‘*-g gene_id*’ option. If a given read was mapped to multiple gene regions, a fraction (1/*x*) of the total number of aligned regions (*x*) were assigned per read per gene (option ‘*-M --fraction*’). QC, mapping, and count statistics were aggregated with MultiQC (Ewels *et al*., 2016). To quantify expression levels for each gene, we calculated transcripts per million (TPM) which normalizes for sequencing depth per sample.

### Differential gene expression analysis

The count dataset from FeatureCounts was imported into R version 4.4.0 (R Core Team, 2024), and normalized using the median of ratios method in the DESeq2 (Love *et al*., 2014) package. This, along with mapping statistics from STAR, revealed one outlier sample which was removed from further analysis. We used principal component analysis and hierarchical clustering methods to quality control at the gene and sample level and explore variation in the data (Fig. S7; Fig. S8). To test for significant differential gene expression between the treatments and life stages in the two experiments across time (“Drought versus Well-Watered”, and “Post-flowering versus Pre-flowering”), we used a likelihood ratio test with the full formula ‘design = ∼ Time + {Treatment, Stage}’ with the reduced formula ‘reduced = ∼ Time’. We also tested significant diurnal gene expression (i.e., the effect of time) for each category (Drought, Watered, Post-Flowering, and Pre-Flowering) separately, with the formula ‘design = ∼ Time’ with the reduced formula ‘reduced = ∼1’. We used ashr (Stephens *et al*., 2023) implemented in DESeq2 to shrink the log2fold change to control for its inflation in smaller values. Our significance threshold was an adjusted (Benjamini-Hochberg) P-value < 0.01, and an absolute value of log2foldchange > 0.58. We used annotated GO IDs to retrieve GO terms related to biological processes using the AnnotationDbi package (Pagès *et al*., 2024) to perform GO enrichment analysis with the enrichr function and gene set enrichment analysis with the GSEA function from the clusterProfiler package v4.12.0 (Wu *et al*., 2021).

### Expression profile clustering and motif analysis

To identify similar expression profiles of differentially expressed (DE) genes between treatment/life stages and across time, we used the degPatterns function in the R package DEGreport (Pantano, 2024) to cluster genes via similarity. For diurnally DE genes that clustered together with core CAM genes, we extracted 2kb up- and downstream to capture regulatory regions (with custom scripts using BEDtools; Quinlan & Hall, 2010). We then used XSTREME (Grant & Bailey, 2021) in the MEME suite (Bailey *et al*., 2015) version 5.5.7 with the default settings, which utilizes several of its own functions to discover (MEME and STREME; Bailey & Elkan, 1994; Bailey, 2021, respectively), annotate (TOMTOM; Tanaka *et al*., 2011), and identify enriched motifs (SEA; Bailey & Grant, 2021) within clusters, generating control sequences from regulatory regions of 1000 randomly-sampled genes. We used the 2022 JASPER CORE DNA non-redundant plant motif database (Castro-Mondragon *et al*., 2021) as the index of known motifs. XSTREME also clusters motifs via similarity (which we will refer to as a “family” of clustered motifs) as its final output. We also ran AME (McLeay & Bailey, 2010) for additional motif enrichment analysis.

## Results

### Genome assembly, annotation, and synteny

The assembled diploid *Cistanthe longiscapa* genome is 647.33 Mbp in size, composed of about 46% repeats (detailed in Table S2), and highly contiguous (L50 = 6 scaffolds, N50 = 44 Mb; 90% of the genome exists in 940 contigs). A total of 28,298 unique protein-coding genes were annotated, which represents a completeness of 97.0% based on BUSCO version v5.7.0 eudicots_odb10.1 database (C: 97.0% [S: 80.4%, D: 16.6%], F: 0.2%, M: 2.8%, n: 2326). The 11 largest scaffolds represent x = 11 chromosomes (2n = 2x = 22 chromosomes; Fig. S1), which is consistent with the signal intensity pattern from the Hi-C contact map (Fig. S3) as well as previous surveys of base chromosome numbers in Montiaceae (Marinho *et al*., 2019). Core CAM genes (Fig. 3a) are scattered across these 11 chromosomes (Fig. S4). Specifically for the *phosphoenolpyruvate carboxylase* (*PEPC*) gene, we found the *1E1a*, *1E1c*, *1E2*, and *2E* copies (Fig. S6), which is consistent with copies found previously in other Montiaceae species (Christin *et al*., 2014).

Comparisons between the *Cistanthe longiscapa* genome and the *Portulaca amilis* genome showed that there are large syntenic blocks that are preserved relative to the evolutionary distance and chromosome number variation, which includes many genes that are in the CAM photosynthetic pathway (Fig. 1e). Furthermore, analyzing the syntenic depth between the two genomes revealed that 37% of the genes in the *C. longiscapa* genome have two homologs in the *Portulaca* genome; inversely, 34 % of *P. amilis* genes have two homologs in the *Cistanthe* genome (Fig. 1d). This reciprocal pattern suggests a whole-genome duplication in a common ancestor, with both genomes losing the duplicated genes at roughly the same rate. This is consistent with a previous study that inferred whole genome duplication at the base of the suborder Portulacineae (Yang *et al*., 2018).

### Induction of CAM phenotypes

We confirmed that our drought treatment does in fact have a physiological effect on the plants by confirming that individuals under drought have a positive carbon assimilation rate at night, which is what we would expect from species using CAM (in 14-hour day length; Fig. 2a). However, we found that *Cistanthe longiscapa* exhibited overall low rates of gas exchange (Fig. 2a). *C. longiscapa* pre-flowering plants that were subjected to drought significantly increased their nighttime acid accumulation (the CAM phenotype) compared to well-watered plants, as measured by the change in titratable acidity between dawn and dusk (Fig. 2b; one-sided Wilcoxon rank sum exact test, p-value = 5.744e^-5^). Moreover, well-watered, post-flowering plants increased their nighttime acid accumulation compared to pre-flowering plants (Fig. 2b; one-sided Wilcoxon rank sum exact test p-value = 0.0603), though the difference in mean ΔH was not as large as the drought versus well-watered comparison from the previous experiment.

**Figure 2.**
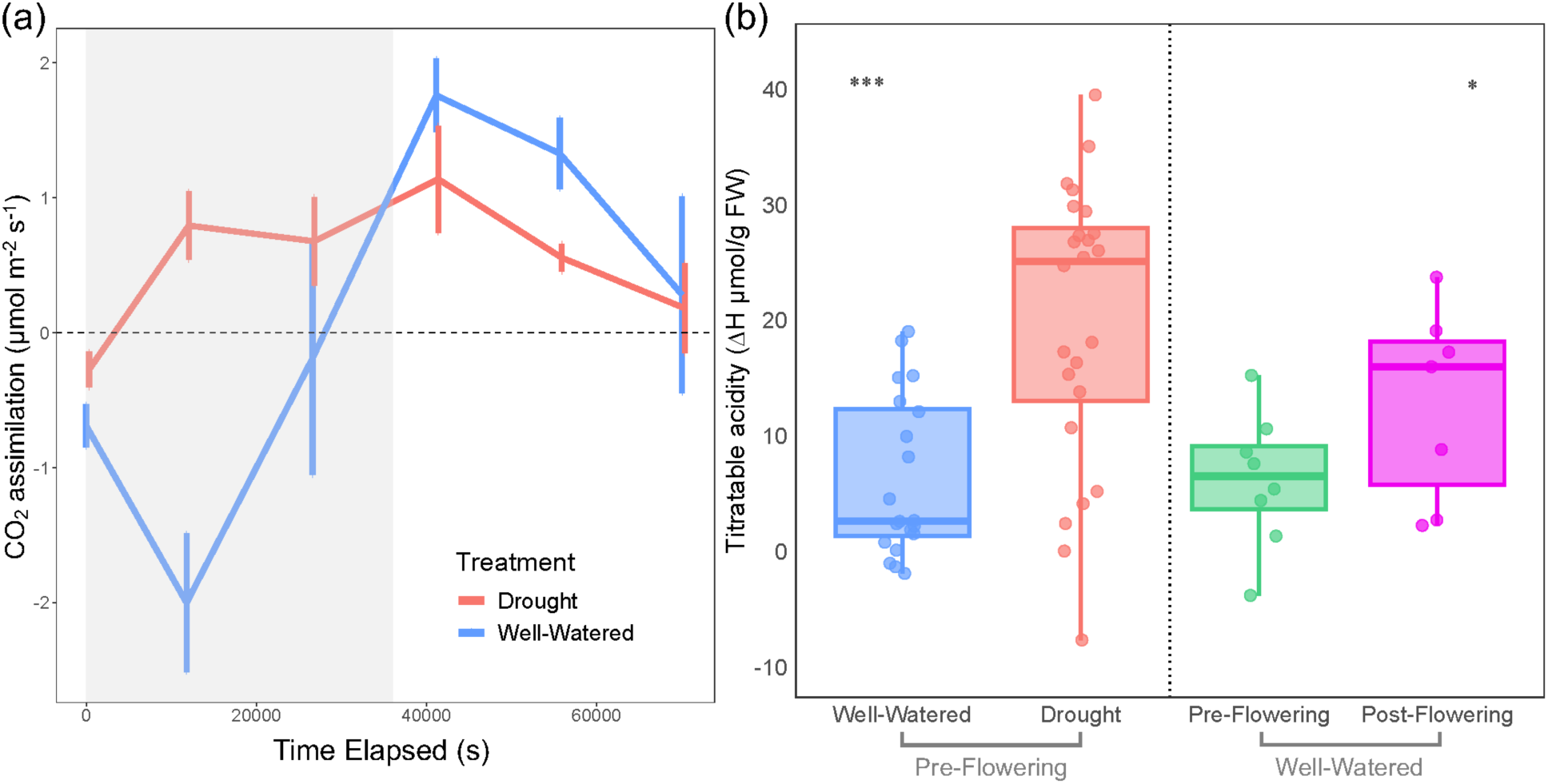
CAM physiology in *Cistanthe longiscapa*. (a) CO_2_ assimilation rate (μmol m^-2^ s^-1^) measured across six timepoints over 24 hours (time elapsed shown in seconds) in an individual under drought (red) and another under a well-watered treatment (blue). Time period with lights off (10 hours) is shaded in gray. Error bars are ± standard deviation taken directly from the LI-6800 machine which was used to measure the assimilation rate. (b) Titratable acidity (in μ equivalents of H^+^; ΔH^+^ µmol g^-1^ ^FW^) assayed among leaf samples of *C. longiscapa* in 1. well-watered treatment and drought treatment (vegetative/pre-flowering individuals), as well as in 2. pre-flowering and post-flowering stages (grown under well-watered conditions). This shows nocturnal acid accumulation (the difference between dawn and dusk collection per sample) which is a measure of CAM activity. Boxplots show median and interquartile range (whiskers = 1.5 × interquartile range). Asterisks indicate significance from Wilcoxon rank sum exact test (“***”, P < 0.001; “*”, P < 0.1).

### Transcriptome assembly and gene abundance

Across 87 RNA libraries, an average of 33.23 million reads per sample were sequenced. We recovered an average of 24.53 million uniquely mapped reads (73.87%) with STAR; 48.21 million aligned sequences (62.72%) were assigned a feature with FeatureCounts (this includes multi-mapped reads from the STAR output). Statistics for individual samples can be found in Dataset S1. A total of 25,504 genes (out of 28,298 annotated genes) were present in the final gene-count dataset. Of these genes, 493 genes were annotated as orthologous copies of the photosynthetic pathway genes (Dataset S2).

In both experiments, the core genes in the CAM carboxylation and decarboxylation pathways (Fig. 3a) showed similar results: *Cistanthe longiscapa* individuals under the drought treatment showed increased abundance and CAM-like expression patterns compared to those under the well-watered treatment, and post-flowering plants showed similar patterns when compared to pre-flowering ones with slightly less overall expression levels of these genes and some exceptions (Fig. 3b,c). We found that the *1E1c* copy of the *PEPC* gene was utilized in the CAM pathway in the drought/well-watered comparison; increased expression of *PPC-1E1c* in early evening in the drought condition is a signature CAM phenotype (Fig. 3b). The post-flowering plants also had CAM-like expression profiles in *PPC-1E1c*, though the highest peak was present right before nighttime rather than early evening; two samples in pre-flowering had spikes of *PPC-1E1c* expression in the night, each at different timepoints (16 & 20 Zeitgeber hours; Fig. 3b), contributing to the large error bars. We identified that the cytosomal *Malate dehydrogenase* gene copy *1-1* (c*NAD-MDH1-1*) and the *malic enzyme* gene copy *4-1* localized in the chloroplast (*cpNADP-ME4-1*) are utilized in the CAM pathway (carboxylation and decarboxylation, respectively; Fig. 3c). *C. longiscapa* using NADP-ME rather than NAD-ME (Fig. S16) for decarboxylation is consistent with the CAM biochemistry seen in *Portulaca* (Gilman *et al*. 2022). Notably, both *phosphoenolpyruvate carboxylase kinase* (*PPCK*) and *phosphoenolpyruvate carboxykinase* (*PCK*) genes were expressed very little in any plant between the two experiments (Fig. S16). This suggests that encoded enzymes are not necessary in the CAM pathway and that PEPC does not need phosphorylation in *C. longiscapa.* This may be due to a mutation in the *C. longiscapa PPC-1E1c* at the position 495 arginine to glutamine (relative to *Portulaca*; Fig. S5), a position where a mutation from arginine to aspartic acid in the *Kalanchoë fedtschenkoi PEPC1* gene has shown to significantly increase PEPC1 activity without phosphorylation (Yang et al., 2017). Lastly, the expression of *pyruvate, phosphate dikinase* (*PPDK*) also followed the overall pattern: increased abundance and CAM-like expression patterns in drought-treated and post-flowering plants compared to well-watered and pre-flowering ones, but with less overall expression levels in post-flowering compared to drought. Normalized abundance plots from all other core genes and gene copies are available in Figure S16.

**Figure 3.**
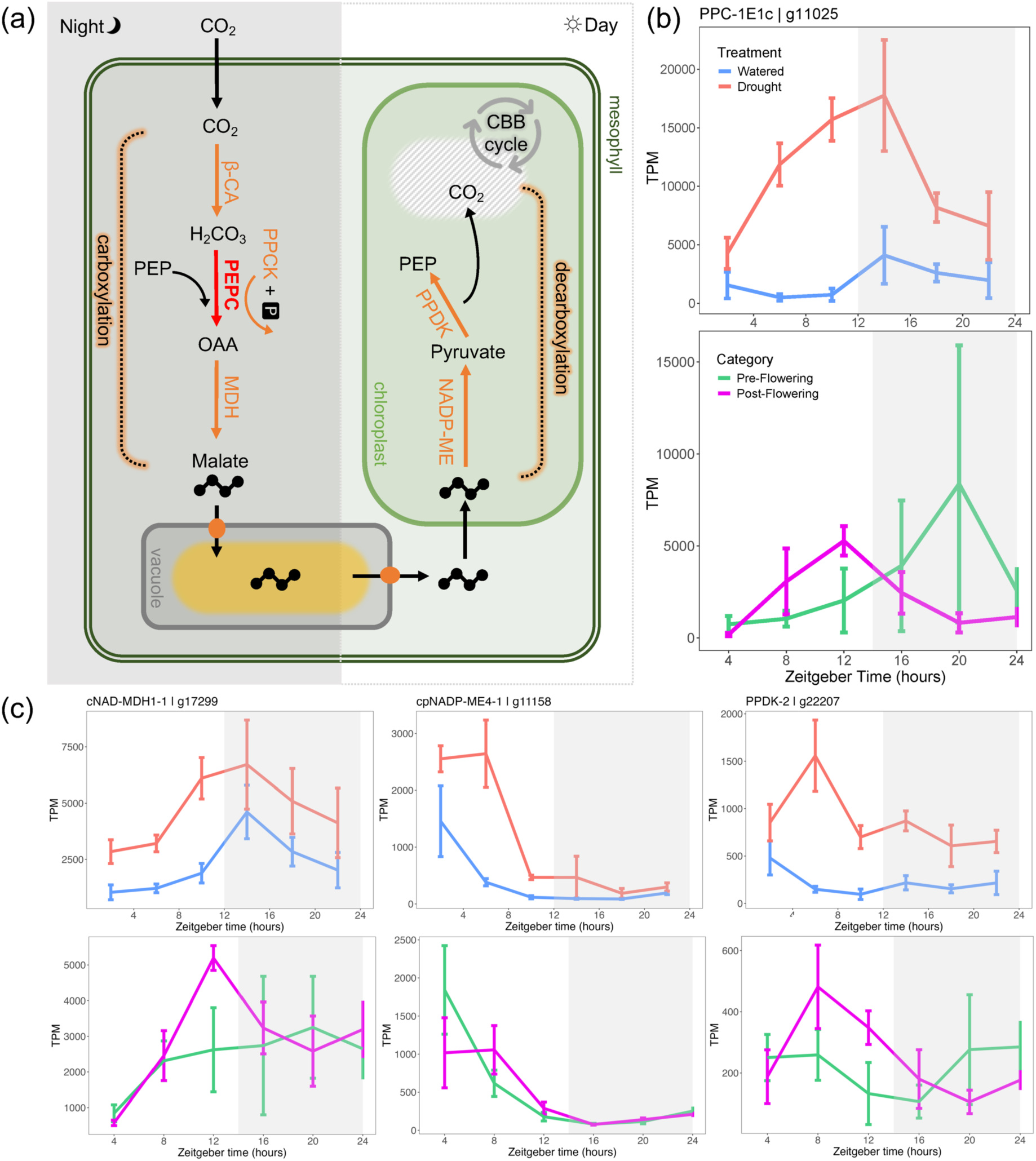
Core CAM gene expression patterns. (a) A simplified schematic of the CAM molecular pathway, showing the nighttime carboxylation and daytime decarboxylation in a mesophyll cell. Core enzymes are shown in orange (except for PEPC, shown in red, for emphasis). β-CA, beta carbonic anhydrase; CBB, Calvin-Bensen-Bassham; MDH, malate dehydrogenase; NADP-ME, NADP-dependent malic enzyme; OAA, oxaloacetate; PPCK, phosphoenolpyruvate carboxylase kinase; PPDK, pyruvate, phosphate dikinase; PEP, Phosphoenolpyruvate; PEPC, PEP carboxylase. (b) Normalized abundance of *PPC-1E1c* (gene g11025 in our annotation) across time (in TPM, transcripts per million). Top panel compares Drought versus Watered, bottom compares Pre-Flowering versus Post-Flowering. Three or four biological replicates per each timepoint; error bars indicate interquartile range. (c) Normalized abundance of *C. longiscapa* copies of *cNAD-MDH* (g17299), cp*NADP-ME* (g11158), and *PPDK* (g22207) genes.

### Differential expression patterns

Many genes showed significant differential expression when tested for the effect of treatment or life stage across time in both experiments (4769 genes in “Drought versus Watered”; 1256 genes in “Post-Flowering versus Pre-Flowering”; Fig. 4a; Fig. S10; Fig. S11; Dataset S3). Several of the top 10 terms in the GO enrichment and GSEA analyses from both experiments were related to photosynthesis, growth, or stress response (Fig. S14). *PPC-1E1c* (p_adj_ = 7.47e^-5^), *cpNADP-ME4-1* (p_adj_ = 1.99e^-4^), and *cNAD-MDH1-1* (p_adj_ = 4.01e^-4^) were differentially expressed (DE) in the “Drought versus Watered” experiment, while other photosynthetic pathway genes were significantly DE in the “Post-Flowering versus Pre-Flowering” experiment (Fig. S18). Around half of the DE genes in the “Post-Flowering versus Pre-Flowering” comparison were shared with those in the “Drought versus Watered” comparison (568 genes; Fig. 4a). Among these 568 shared DE genes, 512 were up- or downregulated in the same direction in both experiments, and only 56 were up- and downregulated in the opposite direction (Dataset S4). Furthermore, among the 568 genes, we found transcription factors and a number of genes that are involved in general stress response, development or reproduction; we highlight these genes in Table 1.

**Figure 4.**
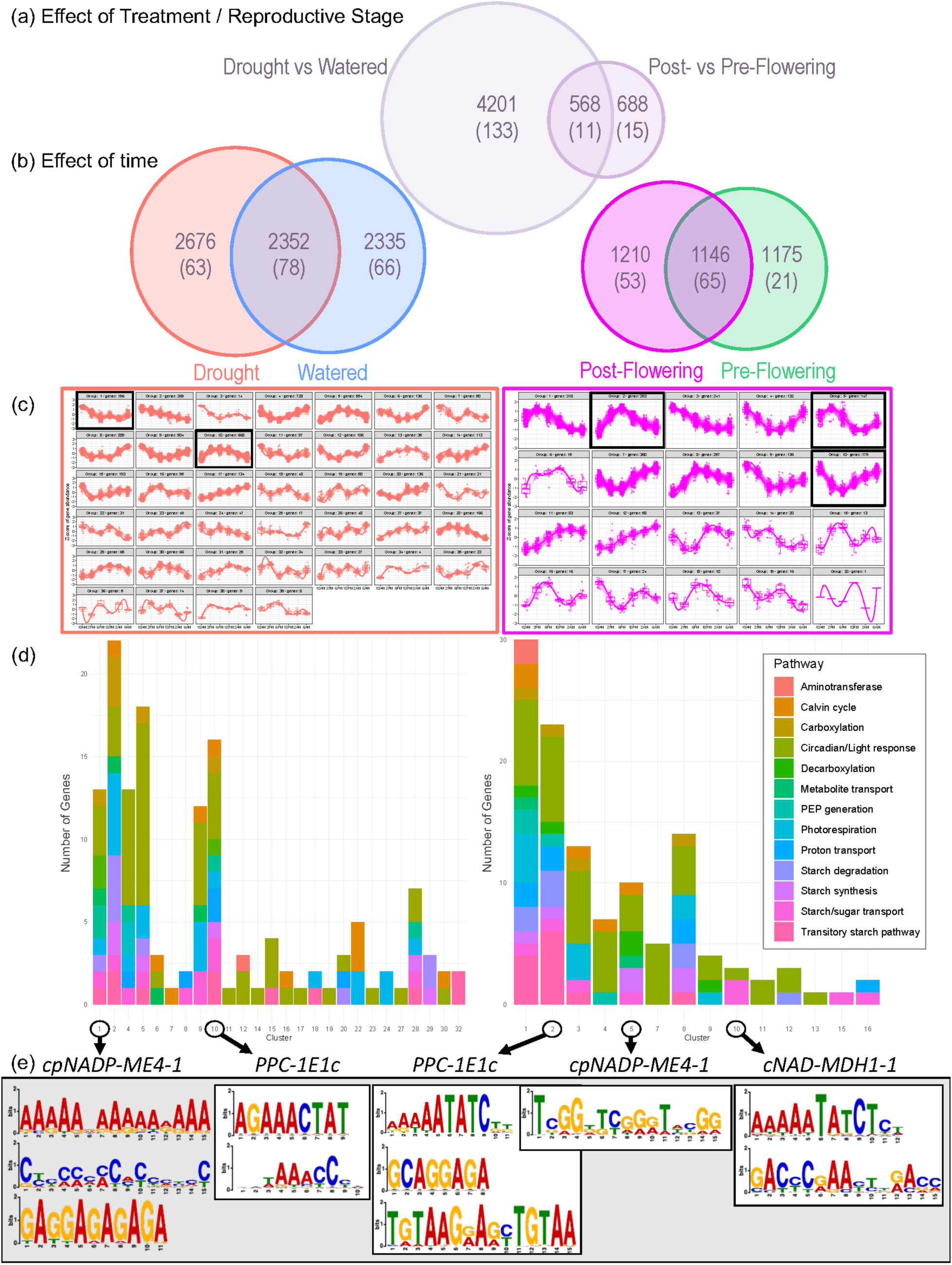
Results from differential gene expression analyses, expression profile clustering, and motif enrichment. (a) Venn diagrams showing the number of differentially expressed genes in the “Drought versus Watered” comparison and “Post-Flowering versus Pre-Flowering” comparisons (testing the effect of treatment or reproductive stage), and the overlap in DE genes between the experiments. Numbers in parentheses represent the number of photosynthetic genes in the differentially expressed sets of genes. (b) Venn diagrams of genes with significant diurnal expression (determined by testing the effect of time) separately in the Drought and Watered treatments and their overlaps, as well as the Post-Flowering and Pre-Flowering stages and their overlaps. Again, numbers in parentheses represent the number of photosynthetic genes in the differentially expressed sets of genes. (c) Clustering of all genes with significant diurnal expression via expression profile similarity in the Drought treatment (left) and Post-Flowering stage (right). Boxplots show median and interquartile range (whiskers = 1.5 × interquartile range) of gene expression in z-score normalized gene abundance; each individual expression profile is shown by thin lines whereas the overall trend is represented with a thick line. Individual cluster plots highlighted with a black box and (d) the number circled correspond to clusters where core CAM genes belong. Bar plots show the number of photosynthetic pathway genes from each corresponding cluster (left panel = Drought; right panel = Post-Flowering). The colors correspond to the photosynthetic pathway identity of each gene, shown in the legend on the right. (e) Enriched motifs in the regulatory regions of all genes (not just the photosynthetic pathway genes) in the clusters where core CAM genes were found; *cNADP-ME4-1* (cluster 1) and *PPC-1E1c* (cluster 10) in the Drought treatment, and *PPC-1E1c* (cluster 2), *cNADP-ME4-1* (cluster 5) and *cNAD-MDH1-1* (cluster 10) in the Post-Flowering stage. Motifs related to DOF zinc finger proteins (“AAAAAAAAAAA”), ERF/DREB family proteins (“CCCCCCCCCCC”), and GAGA-repeats (BBR/BPC class; “GAGGAGAGAGA”) were found to be enriched in all 5 clusters. In all clusters except for Drought cluster 1 (*cNADP-ME4-1*), there were other unique enriched motifs. The only other motif shared is a circadian MYB-related evening element motif (“AAAAATATCTCT”) in cluster 2 (*PPC-1E1c*) and cluster 10 (*cNAD-MDH1-1*) in the Post-Flowering stage.

**Table 1.** Select examples from 568 overlapped differentially expressed genes between the “Drought vs Watered” and “Post-Flowering vs Pre-Flowering” comparisons. **Bold Gene** = transcription factor. Genes related to stress response, development and reproduction are highlighted here, but the full 568 gene list in Supporting Information Dataset S4. “Up” or “Down” in the Regulation column refers to genes being upregulated or downregulated in the Drought treatment and the Post-Flowering stage when compared to the Watered treatment and the Pre-Flowering stage, respectively.

| Gene | Gene IDs | Primary Role | Citation | Regulation |
| --- | --- | --- | --- | --- |
| **AT-hook motif nuclear-localized protein** | g25175 | DNA-binding protein, involved in regulation of growth and development | Zhao *et al.*, 2014 | Up |
| **Auxin Response Factor (ARF)** | g16661 | Regulates auxin-mediated transcription, crucial for plant growth and development | Liscum & Reed, 2002; Van Ha *et al.*, 2013 | Down |
| **B3 domain-containing protein, REM16-like** | g3466 | Involved in development, expressed during early stages of flowering | Mantegazza *et al.*, 2014 | Up |
| **Basic helix-loop-helix (bHLH) leucine zipper transcription factor** | g14561 | Regulates various developmental and metabolic processes | Heim *et al.*, 2003 | Down |
| **Dof zinc finger protein** | g620 | Transcription factor involved in vascular development and photoperiodic flowering | Konishi & Yanagisawa, 2007; Fornara *et al.*, 2009 | Down |
| **Ethylene-responsive transcription factors (ERF/AP2)** | g9235, g19800, g21117 | Binds to Ethylene-Responsive Elements, involved in drought and other stress response | Fujimoto *et al.*, 2000; Sakuma *et al.*, 2002 | Down |
| **Homeobox-leucine zipper proteins** | g4536, g21444, g25444 | Regulates developmental processes, involved in cold stress tolerance, ABA signaling | Henriksson *et al.*, 2005; Cabello *et al.*, 2012; Valdés *et al.*, 2012 | Up* |
| **MADS-box/MEF2 Transcription Factor** | g19958 | Regulates flower development and reproductive processes | Rounsley *et al.*, 1995 | Up |
| **MYB family transcription factors, HRS/PHR-like** | g6103, g14400 | Transcription factors involved in abscisic acid signaling and phosphate starvation response | Wu *et al.*, 2012; Yan *et al.*, 2014; Nagatoshi *et al.*, 2023 | Up (g6103) / Down (g14400) |
| **Squamosa promoter-binding-like proteins** | g2304, g12774, g21618 | Transcription factors regulating flowering and reproductive processes | Klein *et al.*, 1996 | Up (g21618) / Down (g2304, g12774) |
| **TIFY/JAZ family protein** | g6762 | Jasmonate signaling repressors, involved in biotic and abiotic stress responses | Chini *et al.*, 2007, 2017 | Up |
| ABA/WDS Induced Protein | g3570 | Induced by ABA and drought stress, involved in water-deficit stress tolerance | Padmanabhan *et al.*, 1997 | Up |
| Aquaporin Transporters | g13245, g1437 | Water channel proteins, regulate water transport across cell membranes | Maurel & Chrispeels, 2001; Afzal *et al.*, 2016 | Down |
| Cysteine-rich and transmembrane domain-containing protein | g14486 | Involved in stress response/signaling and plant development | Lieber *et al.*, 2011; Xu *et al.*, 2018 | Up |
| DCD Domain-Containing Protein, NRP-like | g16662 | Involved in stress signaling that mediate cell death, associated with ABA signaling | Costa *et al.*, 2008; Zhu *et al.*, 2018 | Up |
| DEAD-BOX ATP-Dependent RNA Helicase 50 | g16819 | Involved in RNA metabolism, splicing, and translation under stress | Li *et al.*, 2023 | Up |
| Late Embryogenesis Abundant (LEA) Proteins | g24748, g11895, g9133 | Seed embryogenesis and drought-stress response and tolerance | Tunnacliffe & Wise, 2007 | Up (g9133),<br>Down (g24748, g11895) |
| Peroxidases | g5240, g5324, g5325, g10678, g17827, | Involved oxidative stress response and stress tolerance | Mittler, 2002 | Down |
| Proline-rich extensin | g16111 | Cell wall structural protein necessary for growth and development | Saha *et al.*, 2013 | Down |
| Small Heat Shock Proteins | g5672, g17836 | Molecular chaperones, protect proteins from aggregation during heat stress | Lindquist & Craig, 1988 | Down |

\*except for g4536 in the Drought vs Watered comparison.

Of the 568 overlapped DE genes between the two experiments, 11 were photosynthetic pathway genes (Fig. 4a; Fig. 5). Six genes related to starch metabolism were all up-regulated in the drought treatment and post- flowering stage, as well as a gene in the photorespiration pathway. Conversely, a gene belonging to the proton transport and another in the circadian/light response pathway were down-regulated in both the drought treatment and the post-flowering stage (compared to the watered treatment and pre-flowering stage, respectively). Two genes were up and downregulated in opposite directions: *Ribulose bisphosphate carboxylase* gene (Calvin cycle) and *Scarecrow-like* gene (circadian/light response).

**Figure 5.**
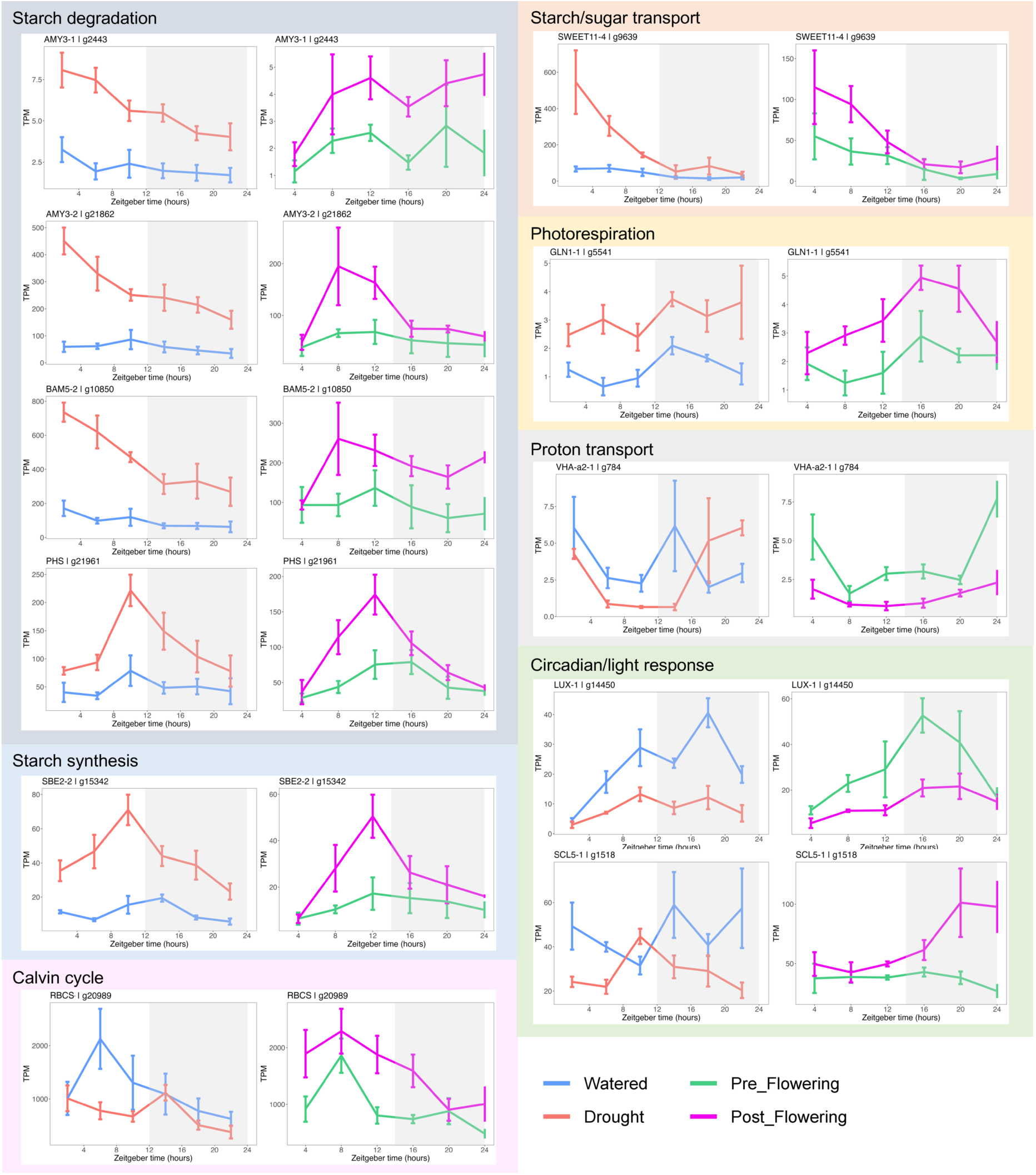
Expression profiles of 11 photosynthesis pathway genes differentially expressed in both experiments. Normalized abundance of these genes are plotted across time (in TPM, transcripts per million), and grouped by photosynthetic pathway identity. Left panels show the results for the Watered versus Drought treatment experiment; right panels show the results for the Pre-Flowering versus Post- Flowering stage experiment. Three or four biological replicates for each treatment x timepoint or stage x timepoint, and error bars indicate interquartile range. AMY, Alpha-amylase; BAM, Beta-amylase; GLN, Glutamine synthetase; LUX, LUX ARRHYTHMO; PHS, Alpha-glucan phosphorylase; RBCS, Ribulose bisphosphate carboxylase small subunit; SBE, 1,4-alpha-glucan-branching enzyme; SCL, Scarecrow-like; SWEET, Bidirectional sugar transporter; VHA, V-type proton ATPase.

Many genes also showed significant differential expression with respect to time (i.e., exhibit diurnal patterns; p_adj_ < 0.01) in each treatment and life stage (5028 genes in the Drought treatment, 4688 in Watered, of which 2352 overlap; 2356 in the Post-Flowering stage, 2321 in Pre-Flowering, of which 1146 overlap; Fig. 4b). *PPC-1E1c* was differentially expressed across time in drought conditions (p_adj_ = 0.0027) and in the post-flowering stage (p_adj_ = 7.49e^-4^). Additionally, *cpNADP-ME4-1* was also diurnally DE in both the drought treatment (p_adj_ = 9.29e^-7^) and post-flowering stage (p_adj_ = 2.11e^-11^), while *cNAD-MDH1-1* was diurnally DE only in the post-flowering stage (p_adj_ = 2.87e^-6^). Various other photosynthetic pathway genes were significantly DE in both the drought treatment and the post-flowering stage, but *not* in the watered treatment or pre-flowering stage; a number of these genes were related to starch metabolism and circadian/light response (Fig. S17). Additionally, photosynthesis GO terms ended up in the top 10 terms from both GO enrichment and GSEA analyses in the drought and post-flowering DE gene set (Fig. S15).

Moreover, we clustered genes via expression profile similarity to look at co-expression of all diurnally DE genes (Fig. 4c; Fig. S11; Dataset S5). In both the drought treatment and post-flowering stage, we found that many photosynthetic pathway genes were co-expressed with each other (in clusters with other genes), though the co-expression could not be explained by pathway identity (Fig. 4d). We also visualized the expression profile clusters for just the DE photosynthetic pathway genes that belong to the “Drought”, “Post-Flowering”, and “Drought versus Watered” sets (141 genes, 118 genes, and 144 genes, respectively; Fig. 4a), which can be found in Figures S12 and S13.

### Motif enrichment

We looked at regulatory regions of all diurnally DE genes that were co-expressed with *PPC-1E1c* in the post-flowering stage (cluster 2; Fig. 4c) and the drought treatment (cluster 10; Fig. 4c) to discover motifs and test for enrichment. Overall, we found: 1) the same circadian rhythm associated motifs enriched in the post-flowering cluster that were enriched in a previous study with genes co-expressed with *PPC-1E1c* in *Portulaca* under drought (Gilman *et al.,* 2022). 2) We found three enriched regulatory motifs shared between the post-flowering and drought clusters that are associated with stress-related gene regulation, plant development, and photoperiodic flower regulation. 3) Between the post-flowering and drought clusters, we found three motifs that are enriched in the post-flowering but *not* in the drought.

Five motif families (from 12 discovered motifs and nine motif matches from the JASPAR CORE dataset) were enriched in the post-flowering cluster (Fig. 4e; Table 2). Outputs from XSTREME and AME were used to confirm that all five motifs are in the regulatory regions of *PPC-1E1c*. One of the enriched discovered motifs (1-AAAAATATCTT; Table 2) belongs to the MYB-related Evening Element (EE) family, associated with transcription factors (TFs) involved in circadian regulation (e.g., RVE8, LHY, CCA; Carré & Kim, 2002). Many plant processes are rhythmic, including those relevant to this study, such as carbon assimilation, stress response, and photoperiodic flowering (McClung, 2019). Another enriched motif (MEME-2; Table 2) belongs to the ERF/DREB family predicted to be involved in stress-related gene expression (Fujimoto *et al*., 2000; Sakuma *et al*., 2002). XSTREME clustered this motif with the also enriched GAGA-repeats (MEME-1, -9, and -5; Table 2; Dataset 6), which are binding sites for BBR/BPC TFs involved in plant development (Sahu *et al*., 2023). Finally, the DOF zinc finger protein-related motifs (MEME-6, Table 2) were enriched, which have been shown to be associated with photoperiodic flowering response regulation in *Arabidopsis* (Fornara *et al*., 2009).

**Table 2.**
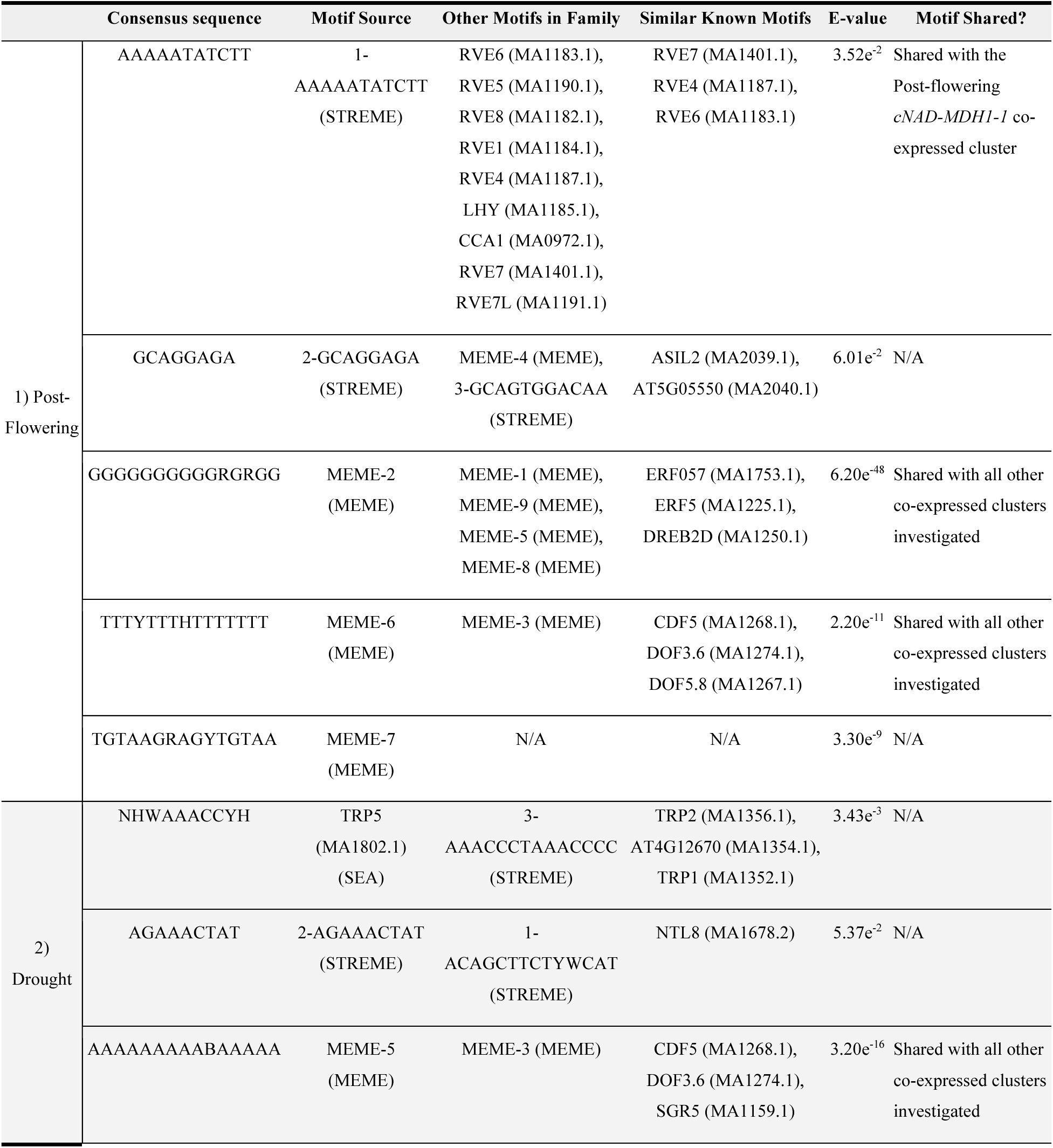
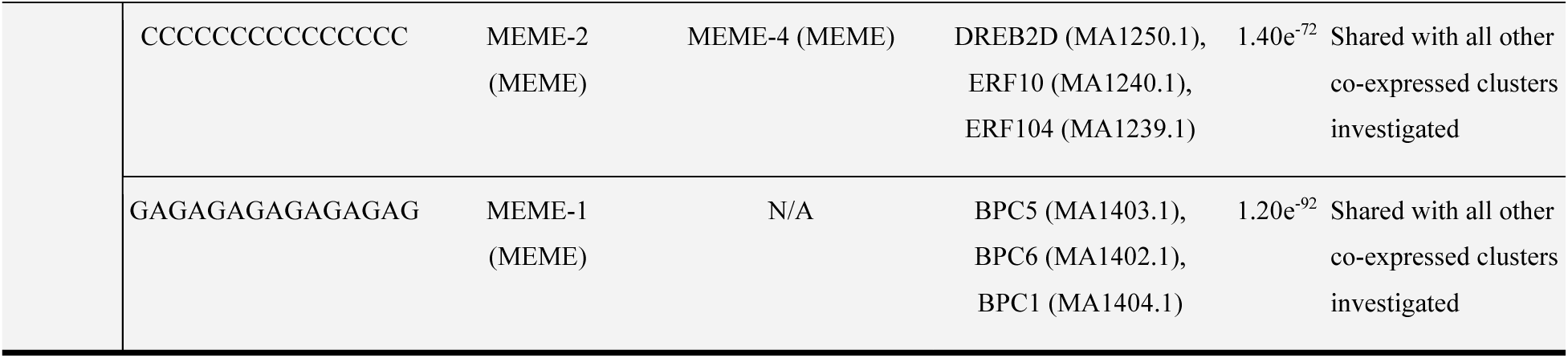
Enriched regulatory motifs from 2 kilobases up- and downstream of co-expressed genes with *PPC-1E1c* from the 1) Post-Flowering stage and 2) Drought treatment experiments. E-values are from the motif discovery step from MEME, STREME, or SEA. Similar known motifs are results from TOMTOM, but only those with similarity E-values of less than 1.0 to the discovered motif are shown. A full result table, including all examined clusters, is available in Dataset S6, and the entire XSTREME output is uploaded on GitHub (https://github.com/anriiam/Cistanthe-RNASeq).

In the drought set, we found five enriched motif families (from 8 discovered motifs and one motif match from the JASPAR CORE dataset), all of which are found in the regulatory regions of the *PPC-1E1c* gene, and three of which were also enriched in the post-flowering cluster (The ERF/DREB family, GAGA, and DOF zinc finger protein-related motifs; Fig. 4e; Table 2). The enriched MYB-related TRP5 motif is associated with TFs primarily involved in telomeric DNA-binding (Hwang *et al*., 2005). Another enriched motif (2-AGAAACTAT; Table 2) is associated with the binding site for the NLT8 (an NAC transcription factor) involved in repressing flowering and germination in *Arabidopsis* (Kim *et al*., 2007, 2008).

We also looked at regulatory regions of all of the diurnally DE genes that were co-expressed with the daytime-expressed CAM pathway genes *cpNADP-ME4-1* in post-flowering (cluster 5) and drought (cluster 1) and *cNAD-MDH1-1* in the post-flowering cluster (cluster 8; Fig. 4c,d,e). In all these clusters, we found the same three motifs enriched (and in the regulatory regions of the genes) as in the *PPC-1E1c* co-expressed clusters from drought and post-flowering, related to gene regulation related to stress (ERF/DREB family), development (GAGA-repeats), and photoperiodic flowering (DOF zinc finger protein-related motifs; Fig. 4e; Dataset S6). Notably, however, there were unique motifs that are only present in the post-flowering clusters but not in the drought. In the post-flowering *cpNADP-ME4-1* cluster, a novel motif was enriched (consensus sequence “TCGGDTCGGRTHCGG”; Dataset S6), which was in the regulatory regions of *cpNADP-ME4-1*. In the *cNAD-MDH1-1* cluster, MYB-related EE motifs (circadian) were enriched (Dataset S6), which were also enriched in the *PPC-1E1c* post-flowering cluster.

## Discussion

### A diversity of CAM behavior in Cistanthe longiscapa

*Cistanthe longiscapa* is endemic to the Atacama Desert and adjacent areas of Chile, one of the most arid landscapes on Earth. Even though they are desert annuals and germinate in response to rainfall, these plants may experience drought for much of development, with effects accumulating over their short lifetime. Because they are facultative CAM plants, if they are experiencing drought around the time they flower and reproduce (i.e., at the end of their lives), they are likely also performing CAM photosynthesis. While CAM is typically considered an adaptation to increase plant water use efficiency, this physiological module may be beneficial for other biological processes as well (Herrera, 2009). Reproduction, particularly flower and seed development, represents a substantial carbon investment (Bazzaz *et al*., 1987), and it is possible that induction of a CAM cycle during reproduction could increase CO_2_ fixation, supplementing daytime C3 CO_2_ fixation with recapturing respiratory CO_2_ or even fixing small amounts of atmospheric CO_2_ at night. Provided a plant is not light-limited and can subsidize the energetic costs of CAM, why would an annual plant with a functional CAM cycle *not* engage it to supplement its carbon uptake during seed production, irrespective of its water status?

Strangely, there are few discussions of possible connections between CAM and reproductive biology in the literature, and direct evidence is lacking. A survey of C3, C4, and CAM plants suggested that CAM plants have the highest flower biomass and seed-to-fruit biomass ratios, and thus higher reproductive carbon costs (but also higher fecundity; Ramírez & Herrera, 2017). In the C3+CAM *Mesembryanthemum crystallinum*, individuals under drought and high-salinity conditions had greater reproductive success, with 10-fold greater seed production, as compared to individuals grown without these conditions (Winter & Ziegler, 1992). This suggests either that greater CAM activity resulted in greater reproductive success, or that when *M. crystallinum* is stressed it increases its reproductive effort (as also reported in *Talinum triangulare*; Taisma & Herrera, 2003). Furthermore, in perennial CAM plant *Dudleya*, flowering individuals displayed higher rates of nocturnal carbon uptake and nighttime acid accumulation as compared to non-flowering individuals, implying an association between reproduction and CAM across clades (Bangal & Lambrecht, 2010).

Our experiments demonstrated that flowering and fruiting plants show elevated nocturnal accumulation of titratable acidity (our phenotypic metric of CAM) relative to non-flowering plants even when under well-watered conditions, though the level of induction was not as significant as their drought response (Fig. 2b). This pattern is also demonstrated in the expression of known CAM genes––CAM response is more strongly upregulated in drought than in post-flowering plants, but many post-flowering gene expression patterns are consistent with a CAM response (Fig 3b, c; Fig 5; Fig S17). Many of the differentially expressed genes shared between the two experiments (“Drought versus Watered”, “Post-Flowering versus Pre-Flowering”) were involved in stress response, plant development, and flowering (Table 1). Moreover, we found that genes co-expressed with nighttime (*PPC-1E1c*) and daytime expressed (*cpNADP-ME4-1*) CAM genes from the post-flowering stage were enriched with regulatory motifs that were unique and absent from their counterparts in the drought treatment (although there were shared motifs related to stress-response, development, and flowering), including a motif related to circadian rhythm (Figure 4e; Table 2; Dataset S6). This suggests that genes involved in the CAM biochemical module have acquired multiple regulatory motifs that are distinct from those involved in drought response and are shared with genes that are upregulated during reproduction.

### A hypothesis for the evolutionary emergence of “reproductive CAM”

Our phenotypic, gene expression, and motif enrichment analyses all tentatively support the delineation of a new kind of C3+CAM phenotype, which we term “reproductive CAM”. It may be possible that this phenotype is widespread in plants that already possess a functional CAM cycle, perhaps particularly in annuals that maximally invest in their reproductive output at the end of their lifecycle. We consider the experiments in *Cistanthe longiscapa* reported here to be preliminary, and encourage further exploration of interactions between CAM expression and reproduction, drought, photoperiod, and temperature. We must also more thoroughly investigate the potential co-regulation of CAM and flowering with finer-scale developmental time-series and more powerful molecular tools (e.g., ATAC-seq or DAP-seq; Buenrostro *et al*., 2013, 2015; Bartlett *et al*., 2017).

In the meantime, it is worth considering a “reproductive CAM” phenotype, and how it might evolve. Phylogenetic analyses strongly support a drought-induced C3+CAM behavior as ancestral to all of Portulacineae (Gilman et al. 2024), suggesting that reproductive CAM is the novel phenotype requiring explanation, rather than a drought-induced CAM evolving from an ancestrally reproductive CAM. It is equally important to highlight the significant caveat that reproductive CAM has not been investigated in other Portulacineae lineages, and so could potentially be widespread. That said, it seems prudent to begin with a model that a drought-induced facultative CAM cycle was ancestrally present and reproductive CAM evolved subsequently. The simplest explanation of reproductive CAM might be that, even in well watered conditions, the simple act of flowering induces some drought stress in individuals, as flowers can often carry significant transpirational burden (Roddy *et al*., 2023). We did not monitor plant water potentials during the experiment and cannot rule out the possibility that flowering and fruiting plants were more water stressed, even under a similar watering regime as vegetative plants. If this were the case, it would be difficult to consider this a new phenotype, as the plant would simply be upregulating CAM in response to drought as it does in other circumstances. Our gene expression data do not really support this model however— there is relatively little overlap in the genes that are co-expressed with CAM genes in the drought and flowering experiments, suggesting that flowering did not simply induce a general “drought-stress module” that happens to include CAM.

Similarly, our experimental design as well as the gene expression data also refute another potential source of correlation: perhaps, reproduction was actually recruited into a more general drought stress response. If so, “reproductive CAM” emerges simply because a switch to flowering is recruited into a drought response, rather than CAM being recruited into a shift to reproduction. But, again, we saw little overlap in drought response and flowering gene co-expression clusters; furthermore, our developmental growth analyses demonstrated very clear and replicated endogenous programming of flowering time (Fig. S2); and additional unpublished work has demonstrated a significant *delay* in flowering in *C. longiscapa* plants grown under chronic stress—the exact opposite of a drought response to initiate flowering (Dauerman *et al*., unpublished).

Instead, it appears that CAM orthologues have acquired additional regulatory motifs (Figure 4e; Table 2; Dataset S6) that connect them to genes involved in reproduction, such that CAM and flowering are at least partially co-regulated because of a direct recruitment of CAM into a reproductive gene regulatory network/networks. We speculate that reproductive CAM could be an example of genetic assimilation (Waddington, 1953; West-Eberhard, 1989), whereby the co-expression of CAM and reproduction is at first due to the facultative drought response and the shared timing of flowering and drought in *Cistanthe longiscapa*’s native environment. Initially, “reproductive CAM” would really be a phenotype that emerges via phenotypic plasticity and a drought-induced CAM response. But the coordinated timing of CAM induction and flowering is adaptive for other reasons as well, facilitating increased productivity and seed set independently of drought stress, and selection would favor maintaining this coordinated timing of expression. Eventually, this emergent new phenotype could become genetically assimilated via recruitment of the CAM pathway into new gene regulatory networks that regulate flowering. The reduced strength of the CAM cycle during flowering as compared to drought, and the mixed expression patterns of CAM genes in the post-flowering stage, suggests that the assimilation and co-option of the full CAM pathway is still incomplete and/or suboptimal in the *C. longiscapa* individuals in our study. It would be well worth surveying other populations in the complicated *C. longiscapa* species complex for variation in the strength and regulation of reproductive CAM.

### Cistanthe longiscapa as a potential “model” C3+CAM species

CAM has evolved in many dozens of lineages independently (Gilman *et al*., 2023) and presents a diverse set of phenotypes that we still know so little about, in terms of their regulation, their potential evolutionary relationships to one another, and even their adaptive significance (Winter *et al*., 2015; Winter, 2019; Edwards, 2023). In the lab, the CAM species best studied at a molecular level are *Mesembryanthemum crystallinum* (Bohnert *et al*., 1988), several *Kalanchoë* (Hartwell *et al*., 2016), and several *Yucca* species (Heyduk *et al*., 2019b) representing only a handful of CAM origins. The recent accumulation of annotated genomes and gene expression studies is opening up new opportunities to study the diverse molecular underpinnings of convergent CAM phenotypes (Cai *et al*., 2015; Ming *et al*., 2015; Brilhaus *et al*., 2016; Wickell *et al*., 2021; Gilman *et al*., 2022; Heyduk *et al*., 2023).

We present *Cistanthe longiscapa* as a new addition to the growing diversity of C3+CAM species available for molecular investigation and laboratory experimentation. Importantly, *C. longiscapa* belongs to an emerging “model clade” (*sensu* Donoghue & Edwards, 2019), the Montiaceae, with increasingly detailed phylogenetic resolution and analyses of organismal evolution that will help contextualize what we learn about its CAM behavior and regulation (Ogburn & Edwards, 2015; Hancock *et al*., 2018, 2019; Chomentowska *et al*., unpublished). The species has recently received attention from ecologists and physiologists (e.g., Stoll *et al*., 2017; Martínez-Harms *et al*., 2022), as one of the most dominant ephemerals in the Atacama desert. This species will readily germinate after sufficient vernalization, is self-fertile, and sets fruit consistently, making it easily tractable to generate multiple generations in the lab. Moreover, we now have a high-quality genome of *C. longiscapa*, which will be important if there is further development of this species as a model system (e.g., establishing transformations). The genome is already key for gene expression studies and comparative genomics—for example, the genomic synteny analyses between *Cistanthe* and *Portulaca* in this study. For the most part, against a backdrop of significant genomic rearrangement, *C. longiscapa* photosynthetic genes belong in syntenic blocks shared with *Portulaca amilis* (Fig. 1e), a relative whose shared common ancestor likely had already established a facultative CAM cycle (Goolsby *et al*., 2018; Gilman *et al*., 2023). A notable exception, of course, was the *1E1c* copy of the *PEPC* gene, which did not stay in its syntenic block with other genes.

### Conclusions

We present *Cistanthe longiscapa* as an excellent new system for understanding the biology and complicated phenotypes of a C3+CAM plant. Most C3+CAM plants are described as “weak constitutive CAM”, showing low levels of CAM expression all the time, or “facultative CAM”, where a CAM cycle is induced (usually by drought) and reversible (Winter, 2019). But if a plant has a functional CAM cycle, there are other possibilities, at least in theory, for its patterns of expression and its recruitment into other aspects of organismal biology, and these are rarely explored. We show here that the CAM cycle is weakly upregulated when a plant is flowering in *C. longiscapa*, and suggest that a novel phenotype, “reproductive CAM”, may be in the initial stages of formation, perhaps via genetic assimilation of an originally drought-induced co-expression of CAM and flowering (Waddington, 1953; West-Eberhard, 1989; Wagner *et al*., 2019). Overall, CAM photosynthesis is an unparalleled system for studies of complex organismal evolution, including but not limited to niche evolution (Smith *et al*., 1986; Edwards & Donoghue, 2006; Bone *et al*., 2015; Hancock *et al*., 2019; Gamisch *et al*., 2021), species diversification (Silvera *et al*., 2009; Arakaki *et al*., 2011; Givnish *et al*., 2014; Bone *et al*., 2015, and many others), genome evolution (Yang *et al*., 2017; Wickell *et al*., 2021), gene regulation and co-expression (Chen *et al*., 2020; Heyduk *et al*., 2022; Gilman *et al*., 2022; Moreno-Villena *et al*., 2022), convergent evolution (Heyduk *et al*., 2019a), and the evolution of trait syndromes and organismal integration (Niechayev *et al*., 2019, Edwards 2023).

## Acknowledgements

We would like to acknowledge and thank Joseph Holtum for providing seeds, and Christopher Bolick and the entire Marsh Botanical Garden and research growth chamber staff for help with plant growth and maintenance. We would also like to thank the Yale Peabody Museum Invertebrate Zoology department for help with microscopy; Ian Gilman, Karolina Heyduk, and Rachel Cohen for advice on analyses, and Patrick Sweeney and Alison Carranza for valuable insights on the project. Funding for this project was provided by the US National Science Foundation Division Of Environmental Biology grant No. 2327957.

## Author Contributions

AC and EJE designed the project. AC grew and harvested plants for genome sequencing and initiated the assembly, which PR completed, scaffolded, and annotated. AC, LW, EJ, SD, VD grew more plants, performed the experiments, and collected data. AC, SD, and VD analyzed the physiological data, and AC and PR analyzed the RNAseq data. AMM and IEP contributed organismal expertise and helped with interpretation of the data. AC, PR and EJE prepared the manuscript, while all authors edited or reviewed and approved the manuscript.

## Data Availability

The genome fasta file and raw RNA sequencing data are available at NCBI under BioProject PRJNA1181828. The annotation file and scripts for genome assembly are available at https://github.com/anriiam/Cistanthe-longiscapa-genome. A script to process RNASeq, raw and normalized RNASeq counts, metadata for RNA samples, and R script for DGE analyses and TPM calculations are available at https://github.com/anriiam/Cistanthe-longiscapa-RNASeq. In the same repository, data and R scripts for visualizing gas exchange and titratable acidity are also available.

## Supporting Information

Dataset S1. RNASeq QC statistics.

Dataset S2. List of photosynthetic gene orthologs.

Dataset S3. Outputs from differential gene expression analyses.

Dataset S4. Overlaps in differentially expressed genes.

Dataset S5. List of genes in co-expressed cluster groups.

Dataset S6. Full list of discovered and enriched motifs.

Figure S1. *Cistanthe longiscapa* chromosome squash image.

Figure S2. *Cistanthe longiscapa* leaf accumulation curves.

Figure S3. *Cistanthe longiscapa* Hi-C contact map.

Figure S4. *Cistanthe longiscapa* genome scaffolds and core CAM gene locations.

Figure S5. *PPC-1E1c* gene sequence comparisons among *Cistanthe longiscapa*, *Portulaca amilis*, and *Portulaca oleraceae*.

Figure S6. *PEPC* gene family tree.

Figure S7. PCA plots from rlog-transformed normalized gene expression.

Figure S8. Heatmaps of rlog-transformed normalized gene expression.

Figure S9. MA plots of mean normalized counts against shrunken log fold change in expression.

Figure S10. Volcano plots from the “Drought versus Watered” treatment comparison and “Post- Flowering versus Pre-Flowering” stage comparison.

Figure S11. Clustering of all significant differentially expressed genes in the two experiments via expression profile similarity.

Figure S12. Clustering of significant differentially expressed photosynthetic pathway genes in the “Drought vs Watered” module and the distribution of pathway identities.

Figure S13. Clustering of photosynthetic pathway genes with significant diurnal expression in the Drought treatment and Post-Flowering stage and the distribution of pathway identities.

Figure S14. Results from GO enrichment analyses among the significant differentially expressed genes and GSEA analyses in both experiments.

Figure S15. Results from GO enrichment analyses among genes with significant diurnal expression and GSEA analyses in the Drought treatment and the Post-Flowering stage.

Figure S16. Expression profiles of other core CAM pathway genes and gene copies from both experiments.

Figure S17. Expression profiles of photosynthetic pathway genes with significant diurnal expression in both Drought treatment and Post-Flowering stage, but not in Watered or Pre-Flowering.

Figure S18. Expression profiles of genes significantly differentially expressed between Post-Flowering and Pre-Flowering stages, but not between Drought and Watered treatments.

Table S1. *Cistanthe longiscapa* leaf samples collected across six time points for RNASeq and corresponding titratable acidity (ΔH^+^) values.

Table S2. Detailed breakdown of repetitive elements of the *Cistanthe longiscapa* genome.

