## Supplemental Information for "A high-quality genome of the mass-blooming desert plant *Cistanthe longiscapa* and its photosynthetic behavior related to drought and life history"

### ***New Phytologist* Supporting Information**

The following Supporting Information is available for this article:

**Dataset S1.** RNASeq QC statistics.

**Dataset S2.** List of photosynthetic gene orthologs.

**Dataset S3.** Outputs from differential gene expression analyses.

**Table S2.** Detailed breakdown of repetitive elements of the *Cistanthe longiscapa* genome.

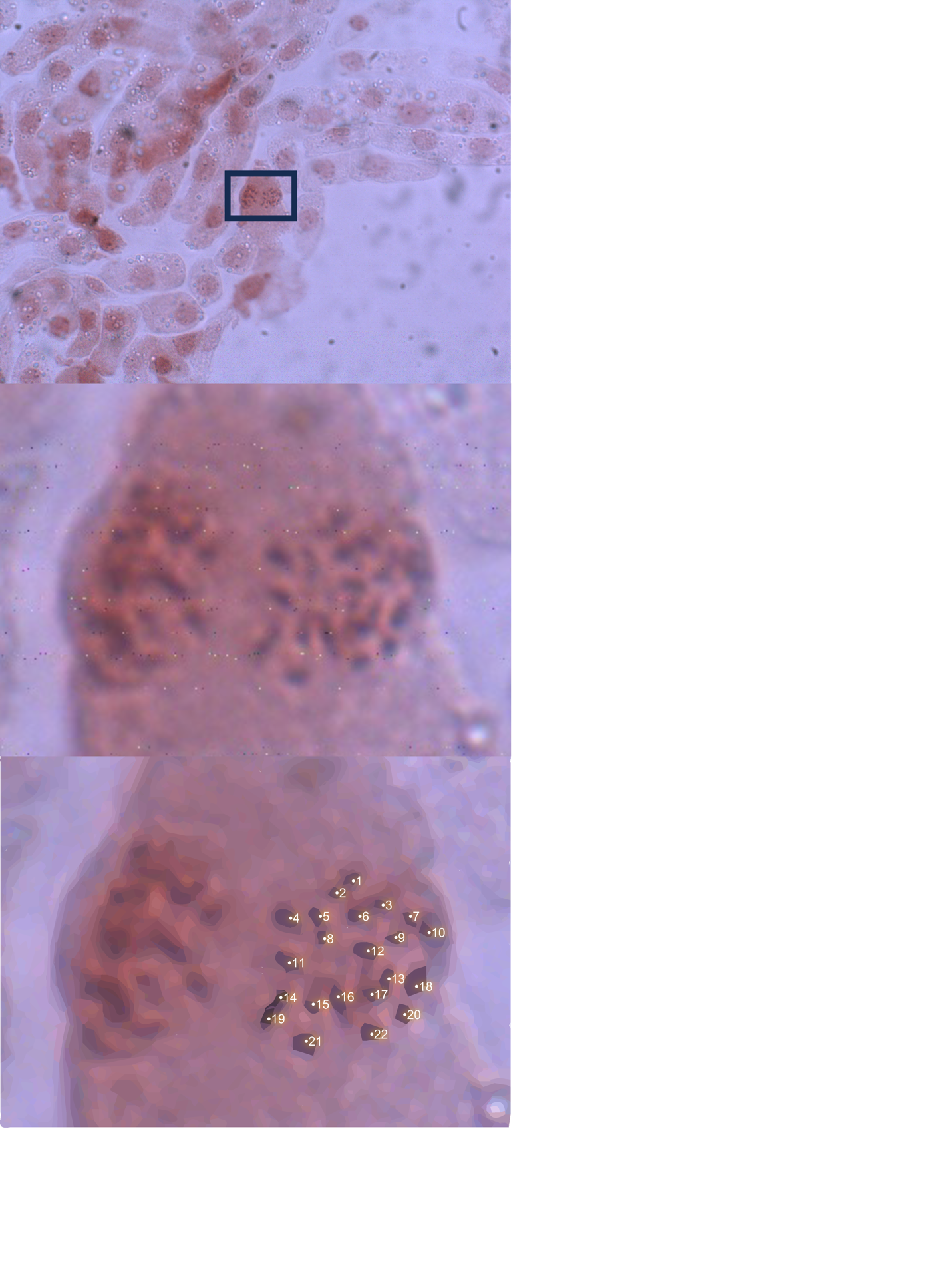

**Figure S1. *Cistanthe longiscapa* chromosome squash image.** Outline of chromosomes were enhanced via Adobe Illustrator® image trace function (bottom panel). 2n = 2x = 22 (x = 11).

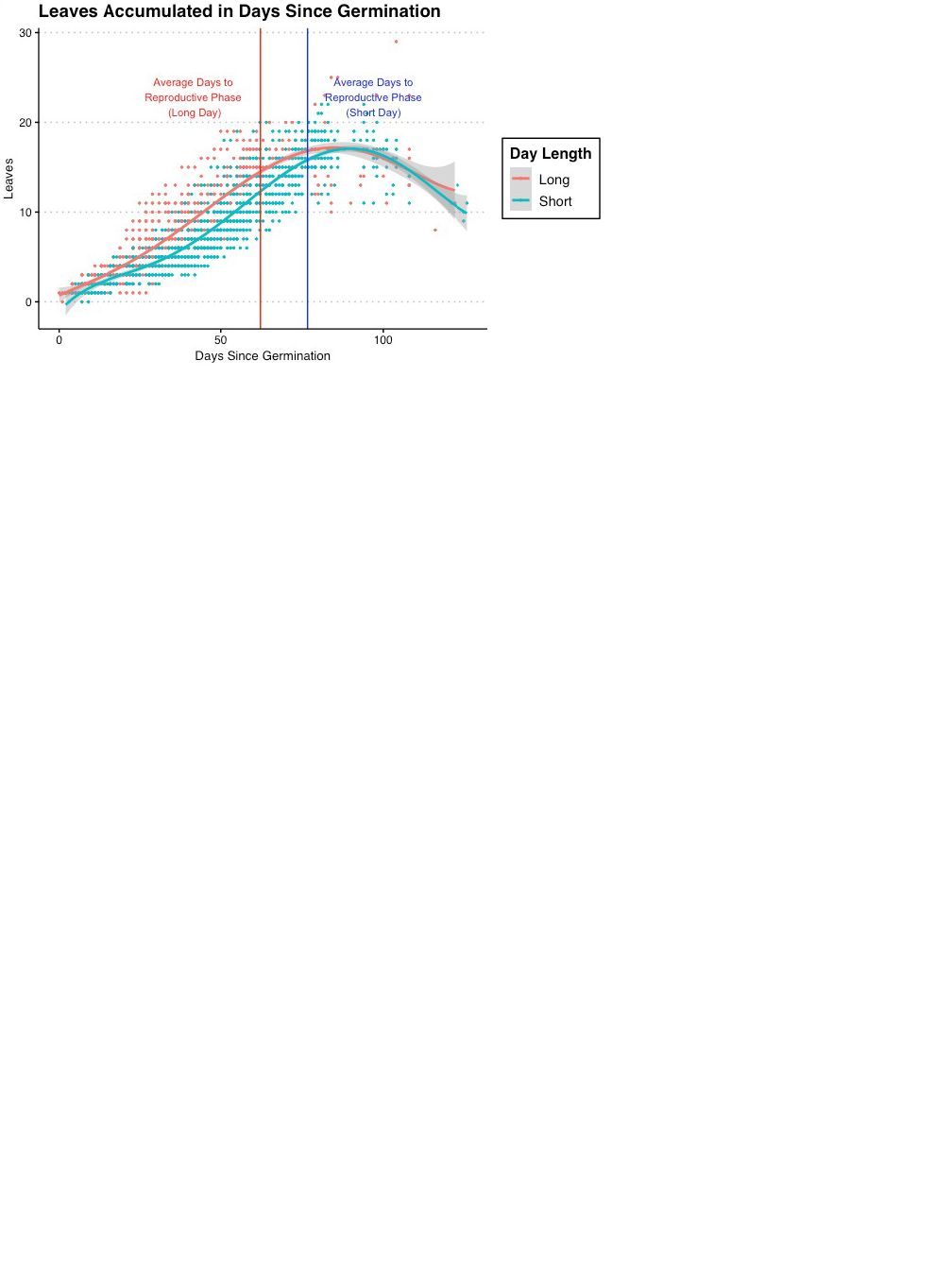

**Figure S2. *Cistanthe longiscapa* leaf accumulation curves.** Red dots represent individuals grown and monitored in long-day (14 hours daylight) chambers, and blue dots represent individuals grown and monitored in short-day (12 hours daylight) chambers. Average number of days to reproductive phase (emergence of a reproductive shoot) is annotated with the vertical lines (62.19 days for long-day; 76.69 days for short-day). The fitted regression curves represent fifth-degree polynomials (gray shading indicates 95% confidence intervals; long-day curve R^2^ = 0.82, short-day R^2^ = 0.84). Both short and long-day plants reach a similar maximum number of leaves around 85 days.

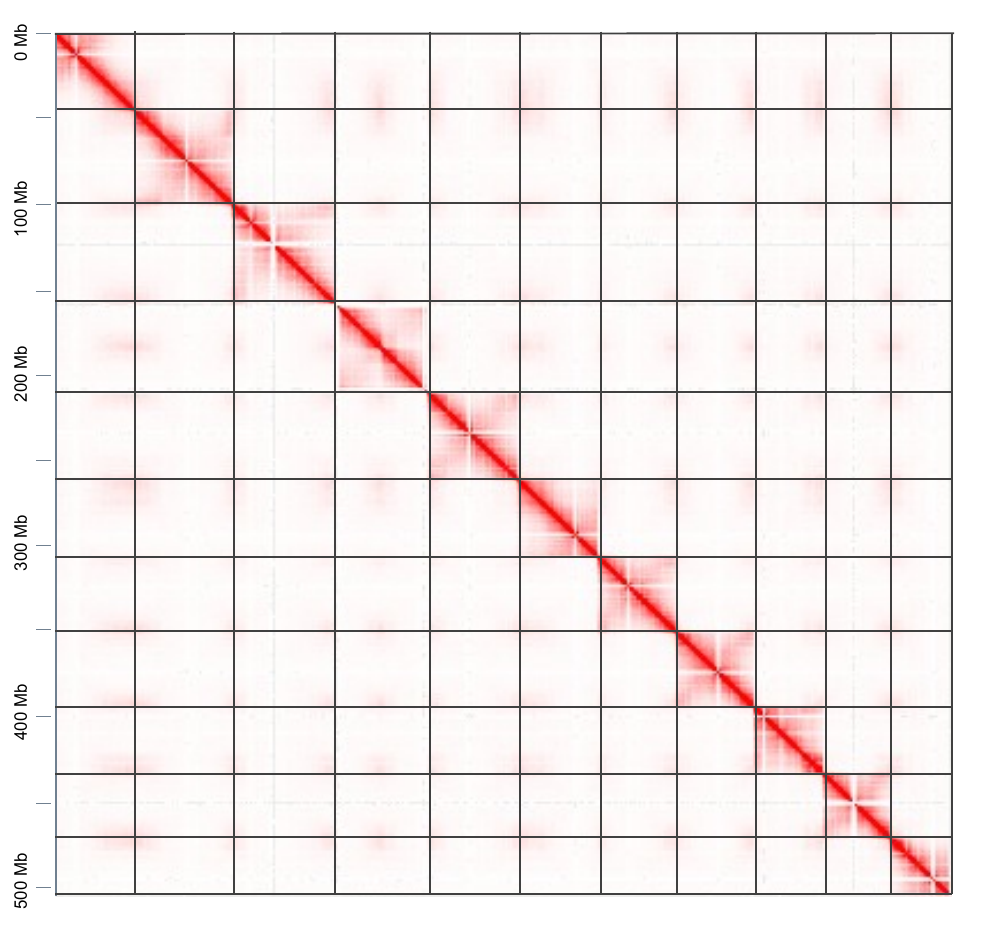
**Figure S3. *Cistanthe longiscapa* Hi-C contact map.** The intensity of signal in red delimits 11 chromosomes.

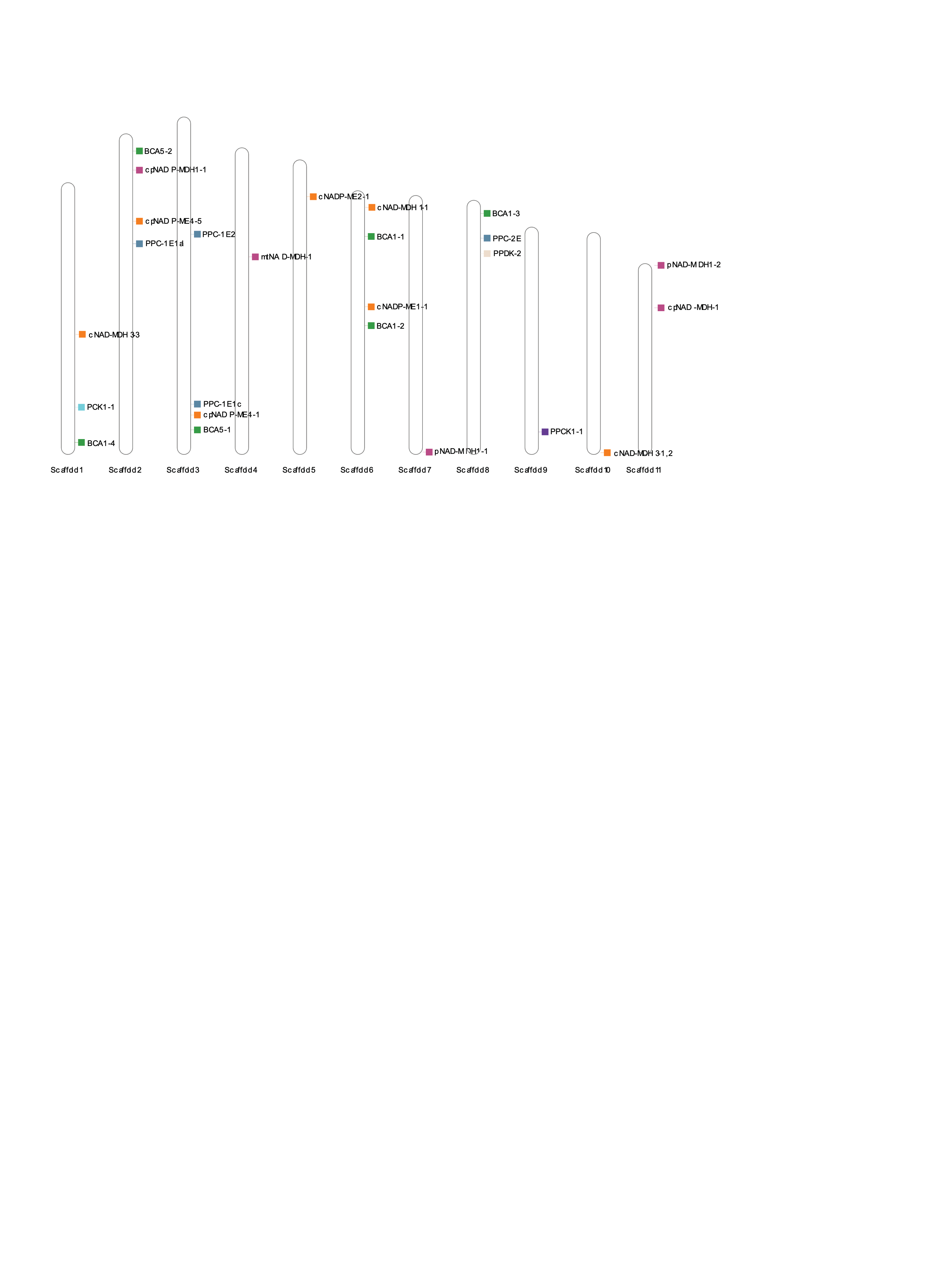

**Figure S4. *Cistanthe longiscapa* genome scaffolds and core CAM gene locations.** The 11 largest scaffolds in the assembly represent the 11 predicted chromosomes. Copies of the core CAM genes are annotated on the scaffolds.

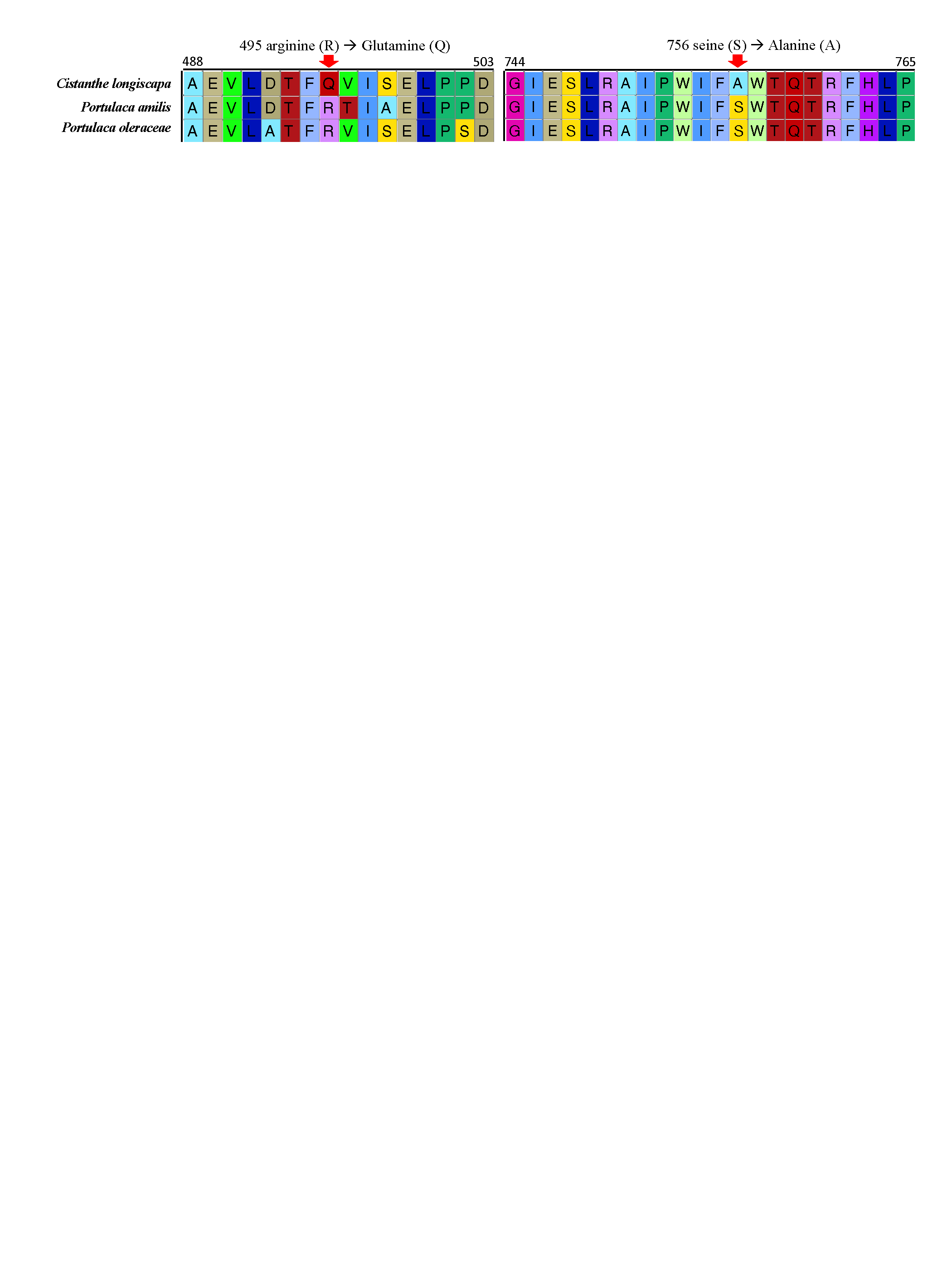

**Figure S5. *PPC-1E1c* gene sequence comparisons among *Cistanthe longiscapa*, *Portulaca amilis*, and *Portulaca oleraceae*.** A snippet of the gene (between positions 488 - 503) is shown to highlight a potentially important mutation in *C. longiscapa* relative to the two *Portulaca* species: 495 arginine (R) → Glutamine (Q).

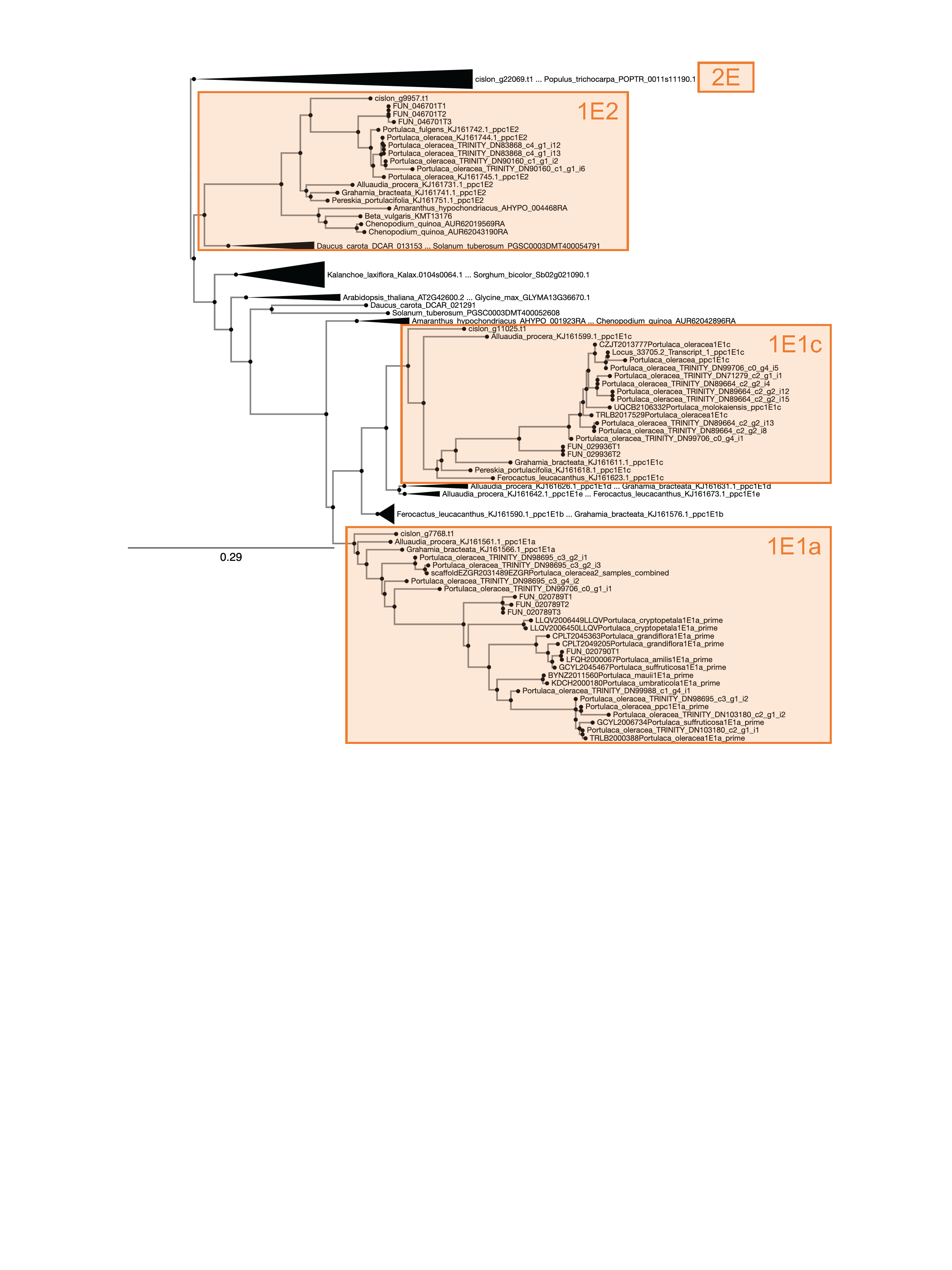

**Figure S6. *PEPC* gene family tree.** Non-important regions of the phylogeny have been collapsed. *Cistanthe longiscapa* copies are labeled with the prefix “cislon” followed by the gene annotation number. In *C. longiscapa,* the *1E1a*, *1E1c*, *1E2*, and *2E* copies were found.

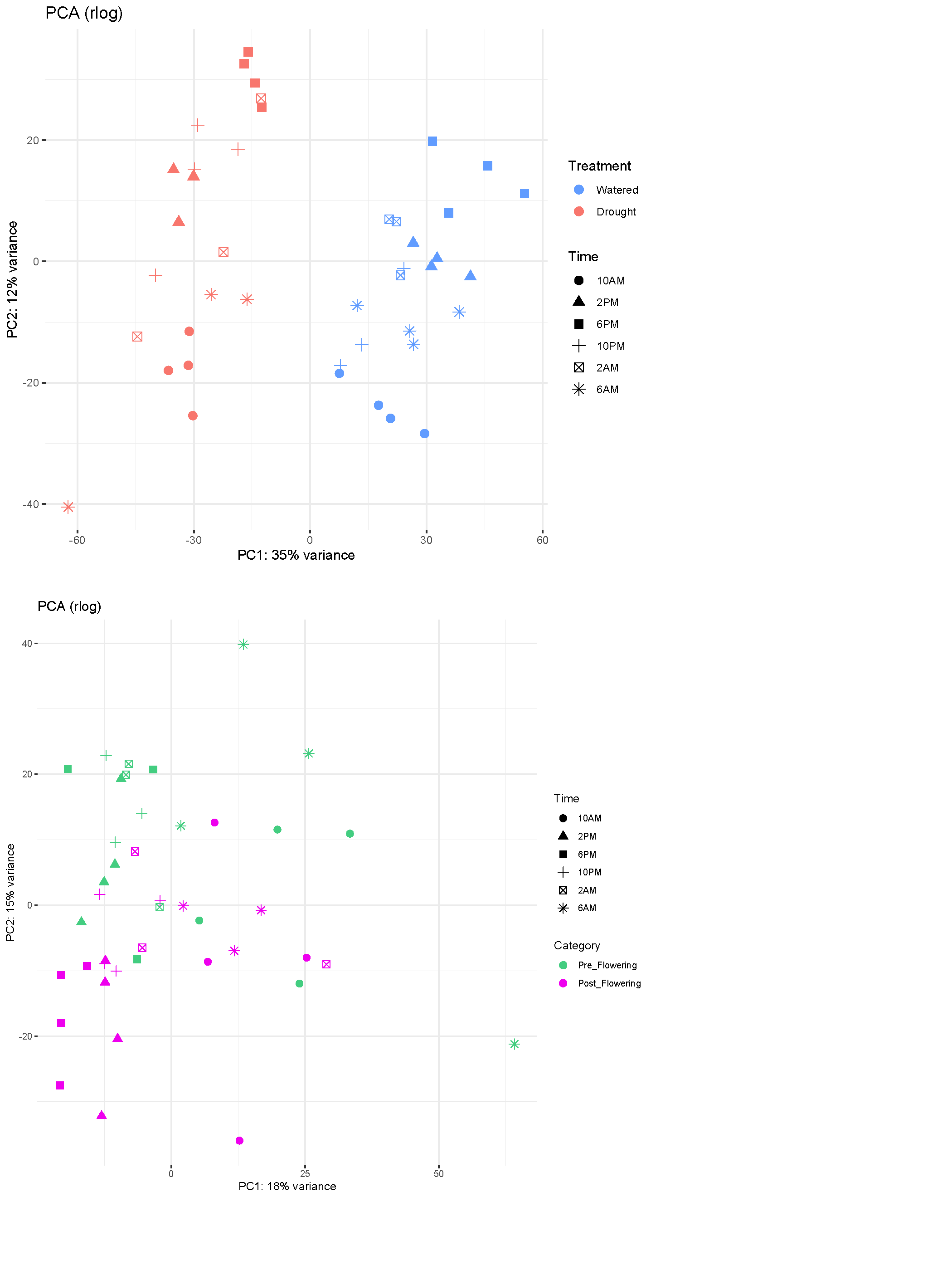

**Figure S7. PCA plots from rlog-transformed normalized gene expression.** Top panel is from the Drought versus Watered treatment comparison experiment; bottom panel is from the Post-Flowering versus Pre-Flowering stage comparison experiment. Each point represents an RNAseq sample.

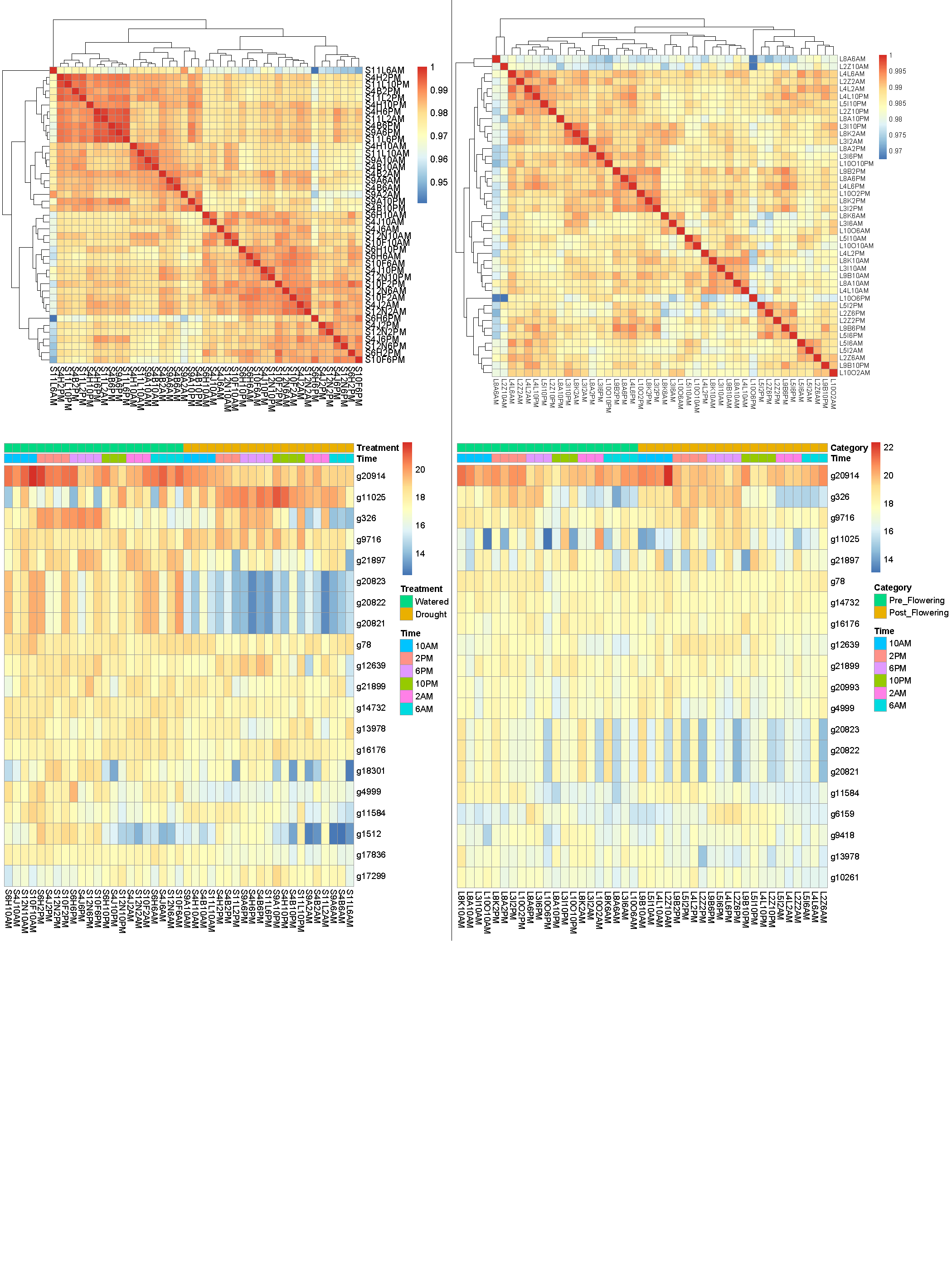

**Figure S8. Heatmaps of rlog-transformed normalized gene expression.** Left panel is from the Drought versus Watered treatment comparison experiment; the right panel is from the Post-Flowering versus Pre-Flowering stage comparison experiment. Top graphs: pairwise correlation between samples (red = higher correlation, blue = lower correlation). Bottom graphs: Expression levels of top 20 most expressed genes across samples (red=higher expression, blue = lower expression).

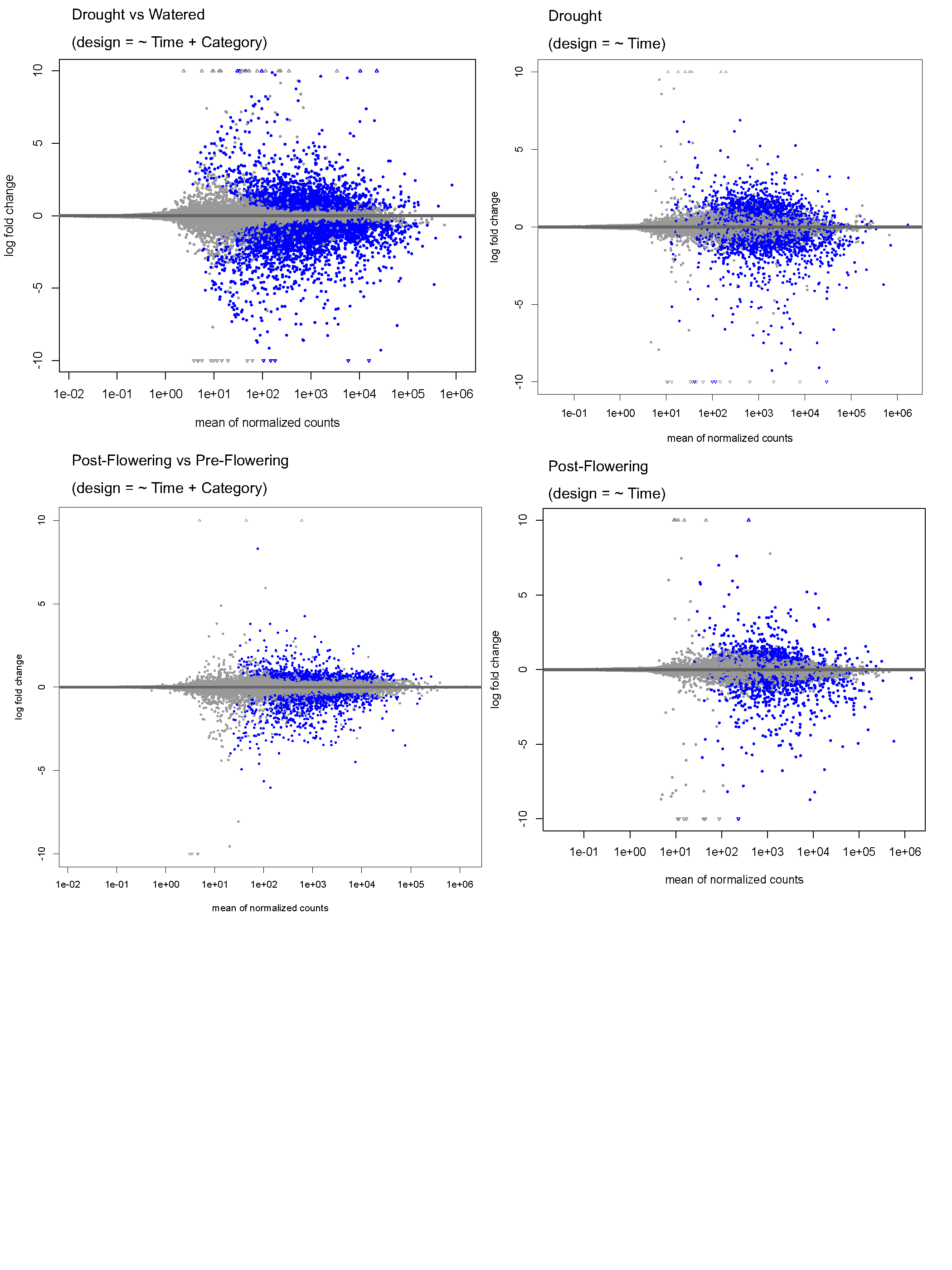

**Figure S9. MA plots of mean normalized counts against shrunken log fold change in expression**. Left two plots are from testing the effect of Drought versus Watered treatments and Post-Flowering and Pre-Flowering stages in DESeq2, and the right two plots are from testing the effect of time in the Drought and Post-Flowering samples in DESeq2. Blue dots = significantly differentially expressed genes (adjusted p-value < 0.01, log fold change > abs{0.58} for left two plots; adjusted p-value < 0.01 for right two plots).

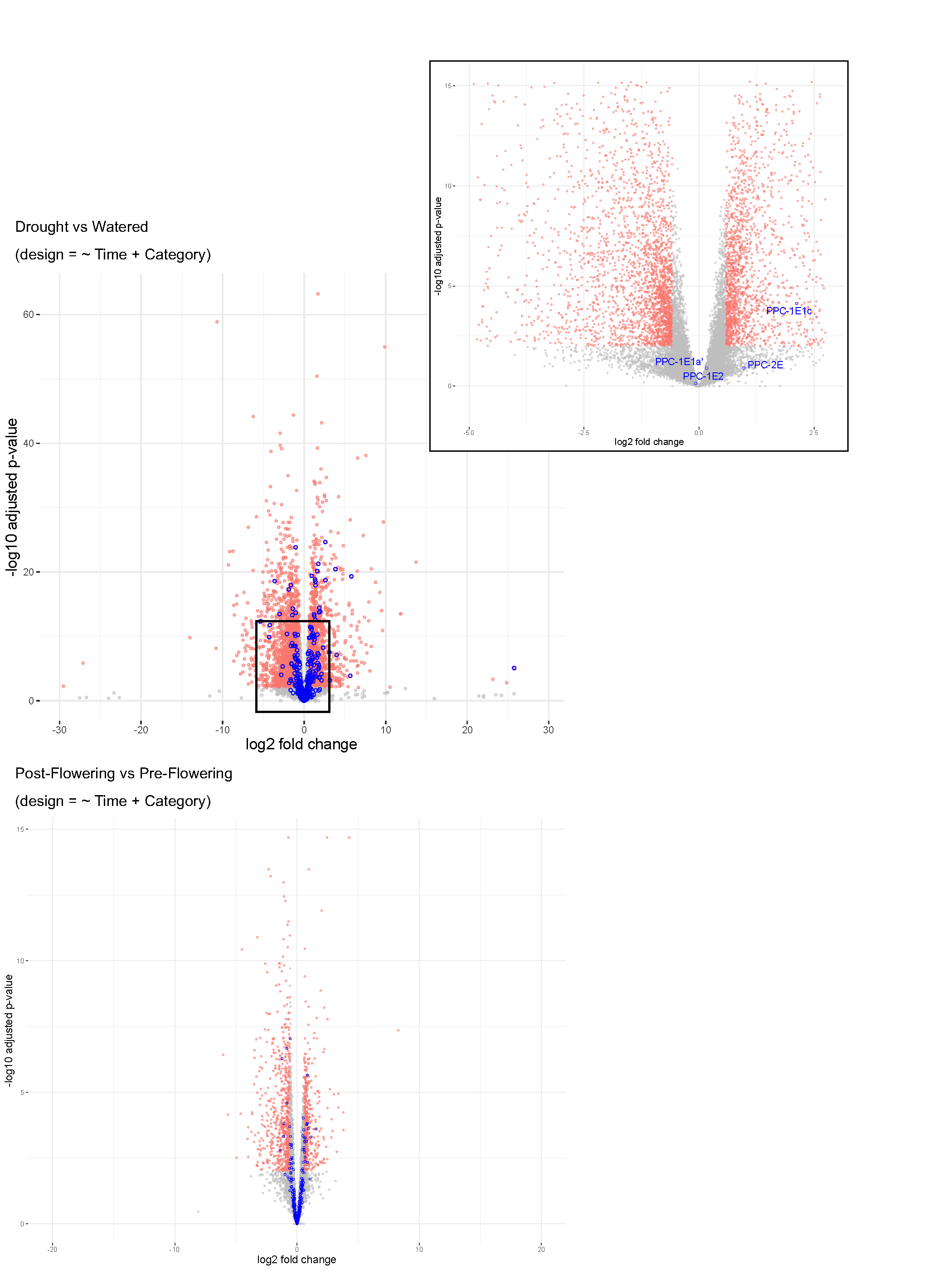

**Figure S10. Volcano plots from the “Drought versus Watered” treatment comparison and “Post-Flowering versus Pre-Flowering” stage comparison.** Red dots = significantly differentially expressed genes (adjusted p-value < 0.01, log fold change > abs{0.58}), blue dots = photosynthetic pathway genes. Location of *PEPC* gene copies from the former experiment are shown in the zoomed panel.

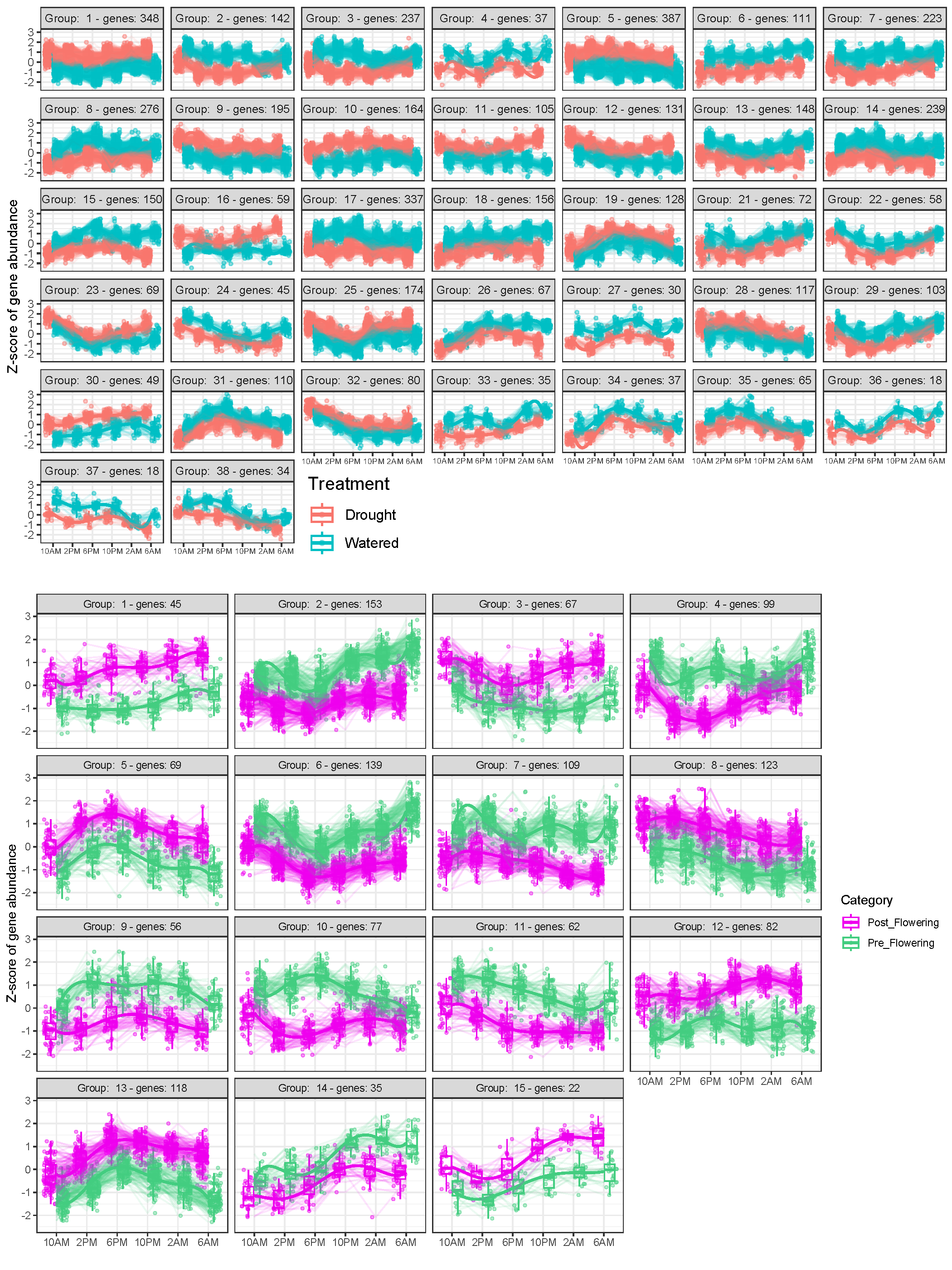

**Figure S11. Clustering of all significant differentially expressed genes in the two experiments via expression profile similarity.** Top panel is from the “Drought vs Watered” DE module, and bottom panel is from the “Post-Flowering versus Pre-Flowering” DE module. Boxplots show median and interquartile range (whiskers = 1.5 × interquartile range) of gene expression in z-score normalized gene abundance; each individual sample expression profile is shown by thin lines whereas the overall trend is represented with a thick line.

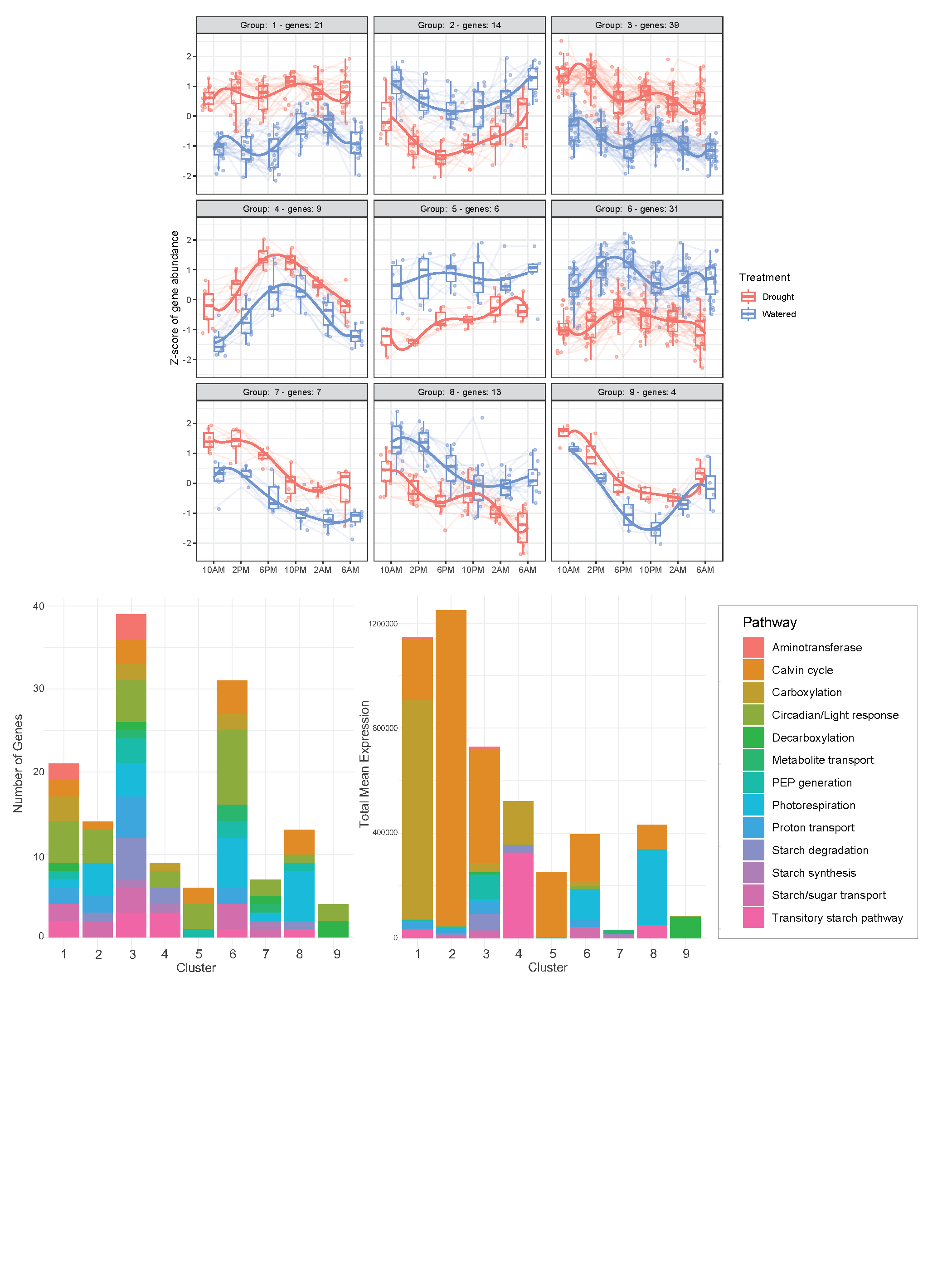

**Figure S12. Clustering of significant differentially expressed photosynthetic pathway genes in the “Drought vs Watered” module and the distribution of pathway identities.** 133 photosynthetic pathway genes in the significantly DE module in the “Drought vs Watered” treatment comparison were clustered via expression profile similarity. Boxplots show median and interquartile range (whiskers = 1.5 × interquartile range) of gene expression in z-score normalized gene abundance; each individual sample expression profile is shown by thin lines whereas the overall trend is represented with a thick line. Bar plots show the number of genes (bottom left) and the total mean expression of genes (bottom right) from each corresponding cluster; the colors correspond to the photosynthetic pathway identity of each gene, shown in the legend on the right.

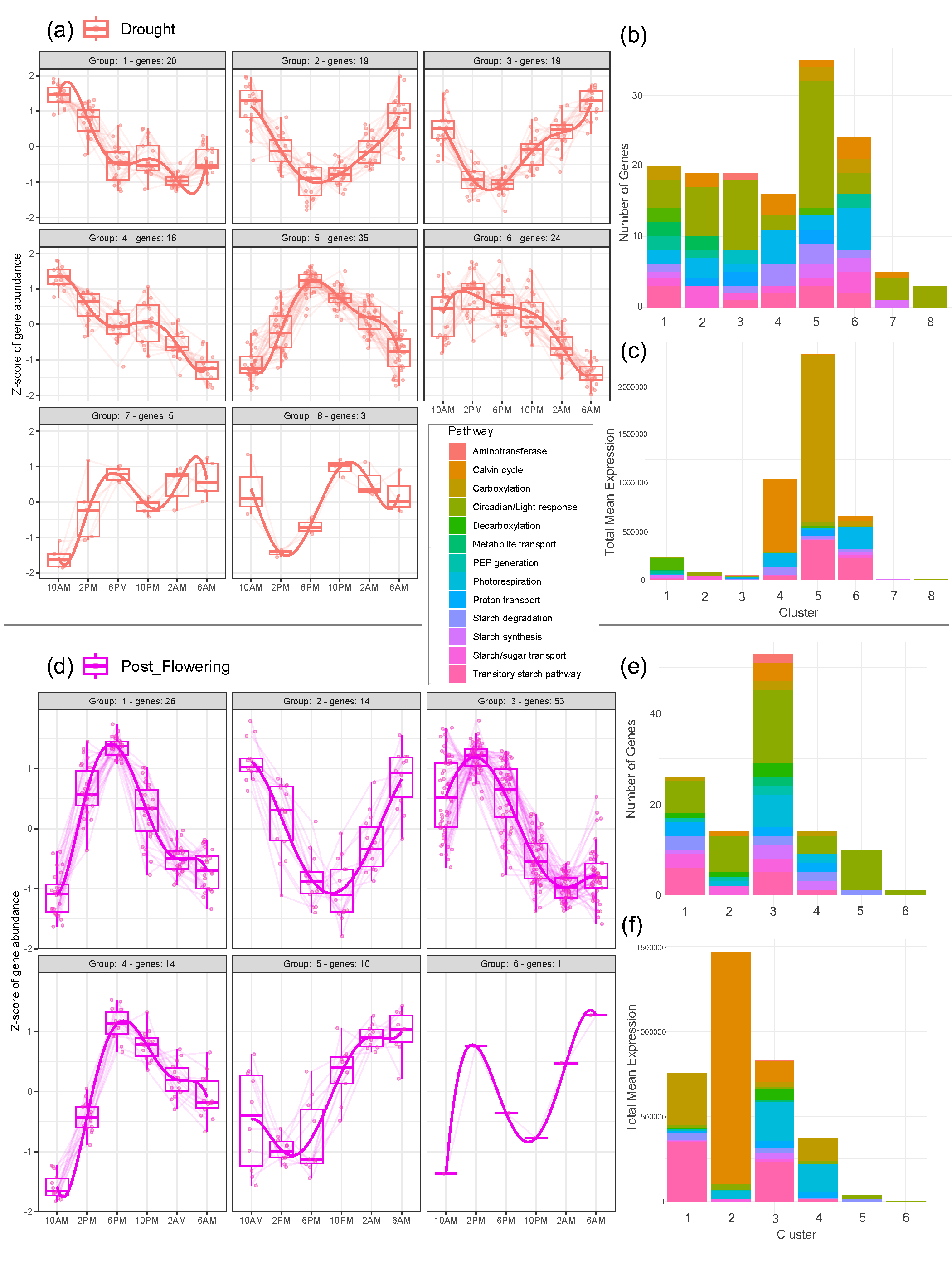

**Figure S13. Clustering of photosynthetic pathway genes with significant diurnal expression in the Drought treatment and Post-Flowering stage and the distribution of pathway identities.** 141 and 118 photosynthetic pathway genes in the (a) Drought and (b) Post-Flowering significantly DE gene sets, respectively, were clustered via expression profile similarity. Boxplots show median and interquartile range (whiskers = 1.5 × interquartile range) of gene expression in z-score normalized gene abundance; each individual expression profile is shown by thin lines whereas the overall trend is represented with a thick line. Bar plots show (b, e) the number of genes and (c, f) the total mean expression of genes from each corresponding cluster; the colors correspond to the photosynthetic pathway identity of each gene.

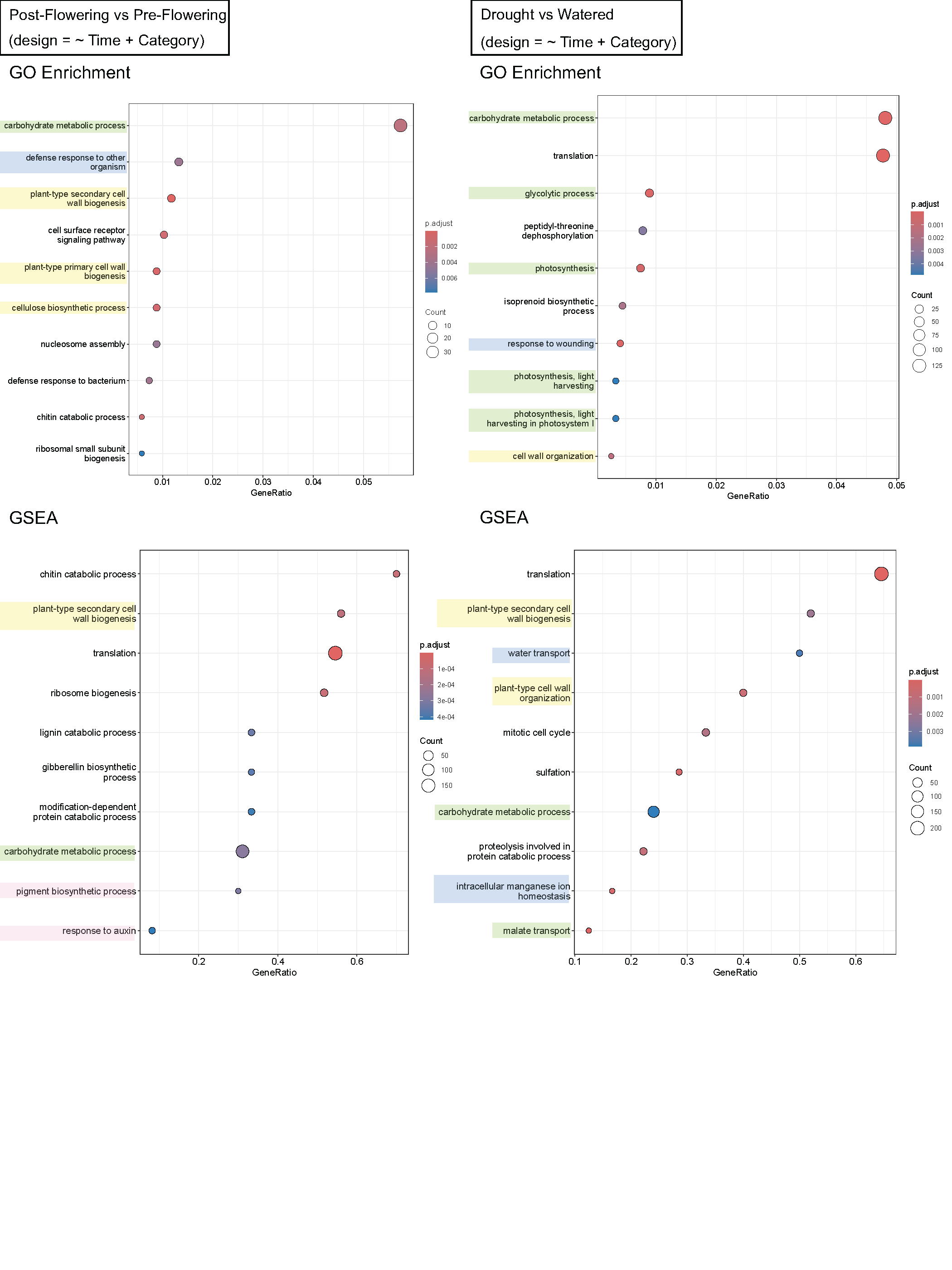

**Figure S14. Results from GO enrichment analyses among the significant differentially expressed genes and GSEA analyses in both experiments.** Top 10 “Biological Process” GO terms (ranked by gene ratio) are shown for each analysis. GO terms related to CAM or photosynthesis (green), plant growth (yellow), stress response (blue), and flowering/reproduction (pink) are highlighted. Gene ratio (the x-axis) represents the proportion of genes associated with the GO term relative to the total number of genes in that category.

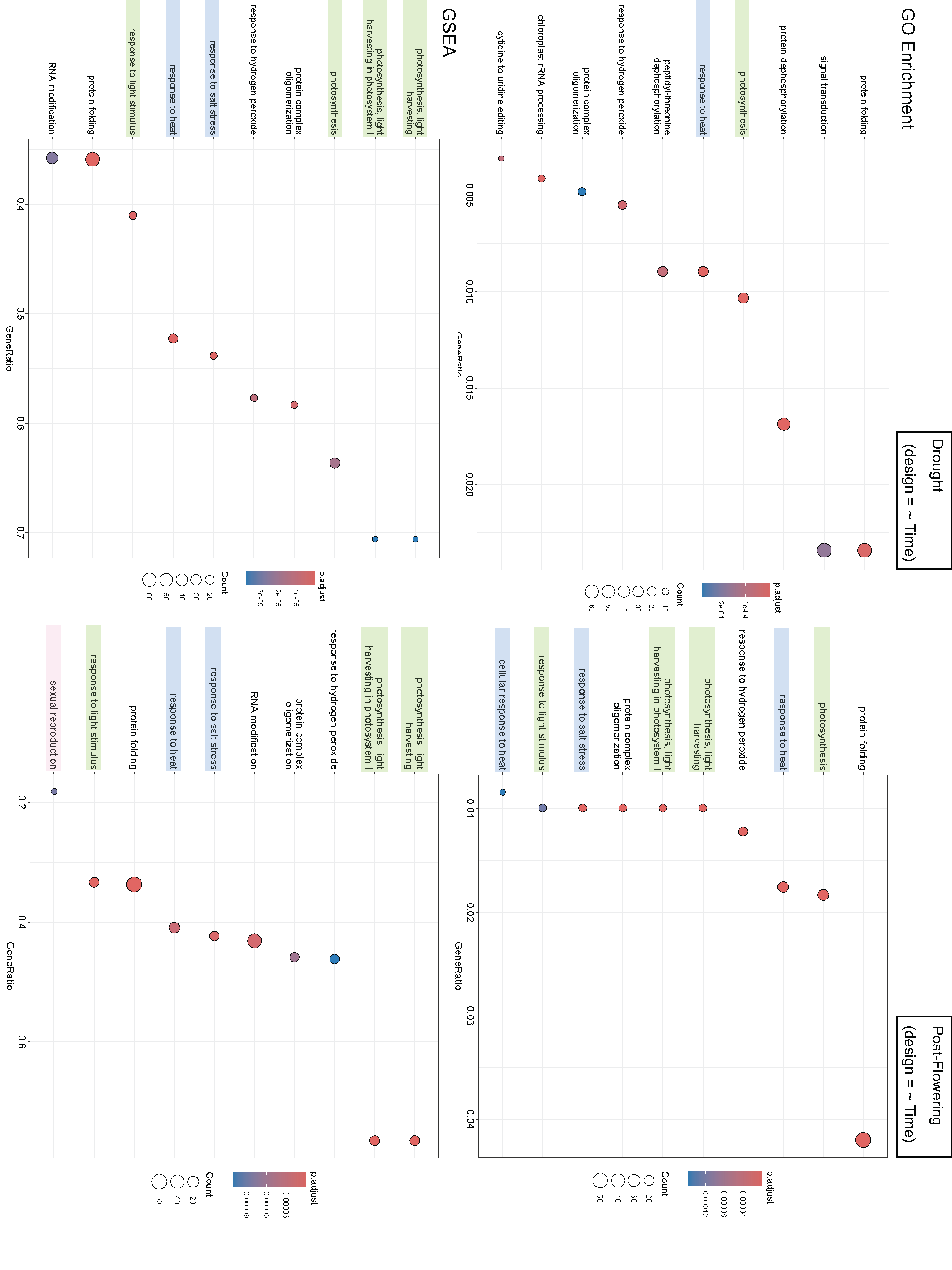

**Figure S15. Results from GO enrichment analyses among genes with significant diurnal expression and GSEA analyses in the Drought treatment and the Post-Flowering stage.** Top 10 “Biological Process” GO terms (ranked by gene ratio) are shown for each analysis. GO terms related to CAM or photosynthesis (green), stress response (blue), and flowering/reproduction (pink) are highlighted. Gene ratio (the x-axis) represents the proportion of genes associated with the GO term relative to the total number of genes in that category.

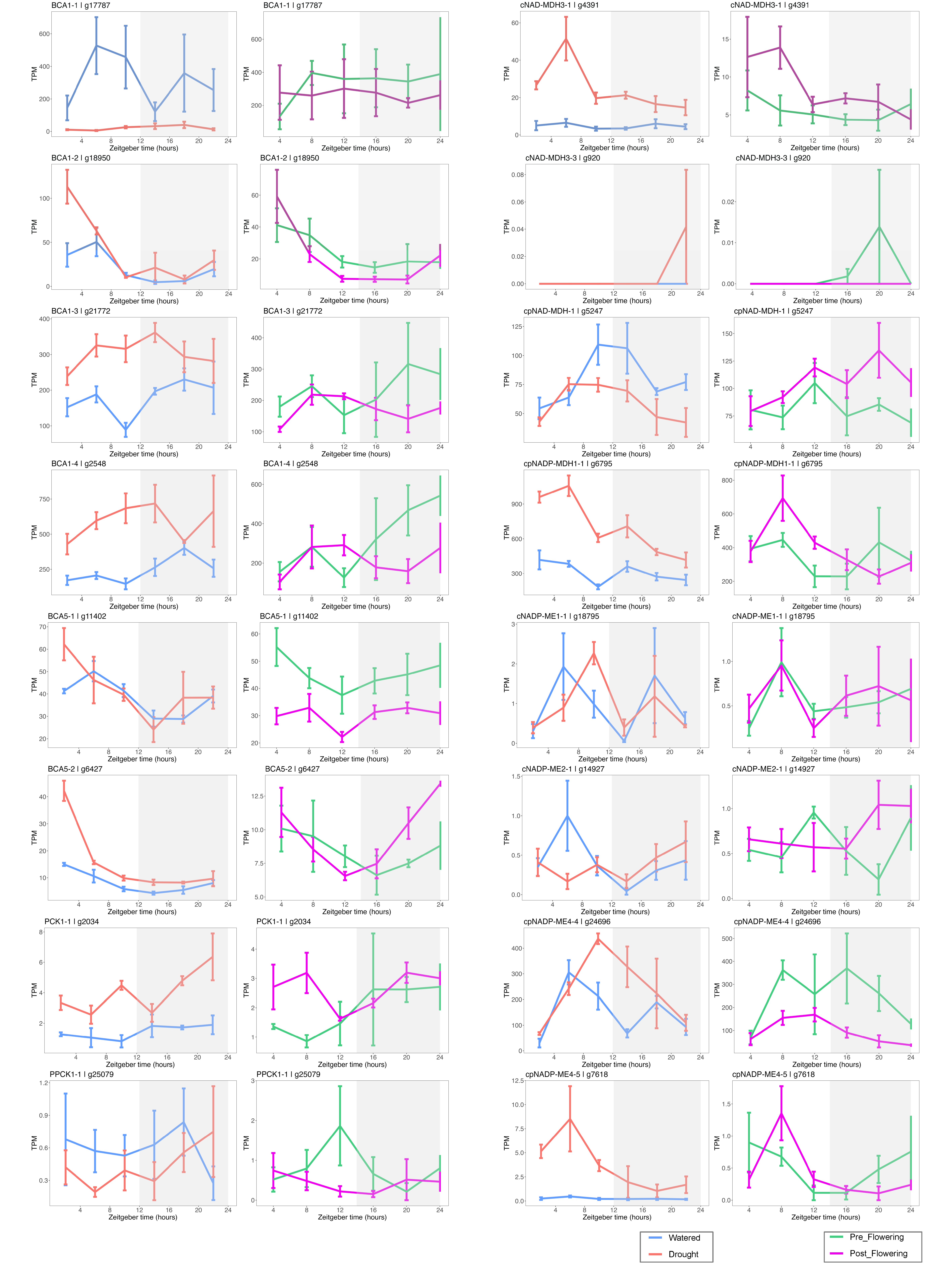

**Figure S16. Expression profiles of other core CAM pathway genes and gene copies from both experiments.** Normalized abundance of these genes are plotted across time (in TPM, transcripts per million). Three or four biological replicates at each timepoint; error bars indicate interquartile range.

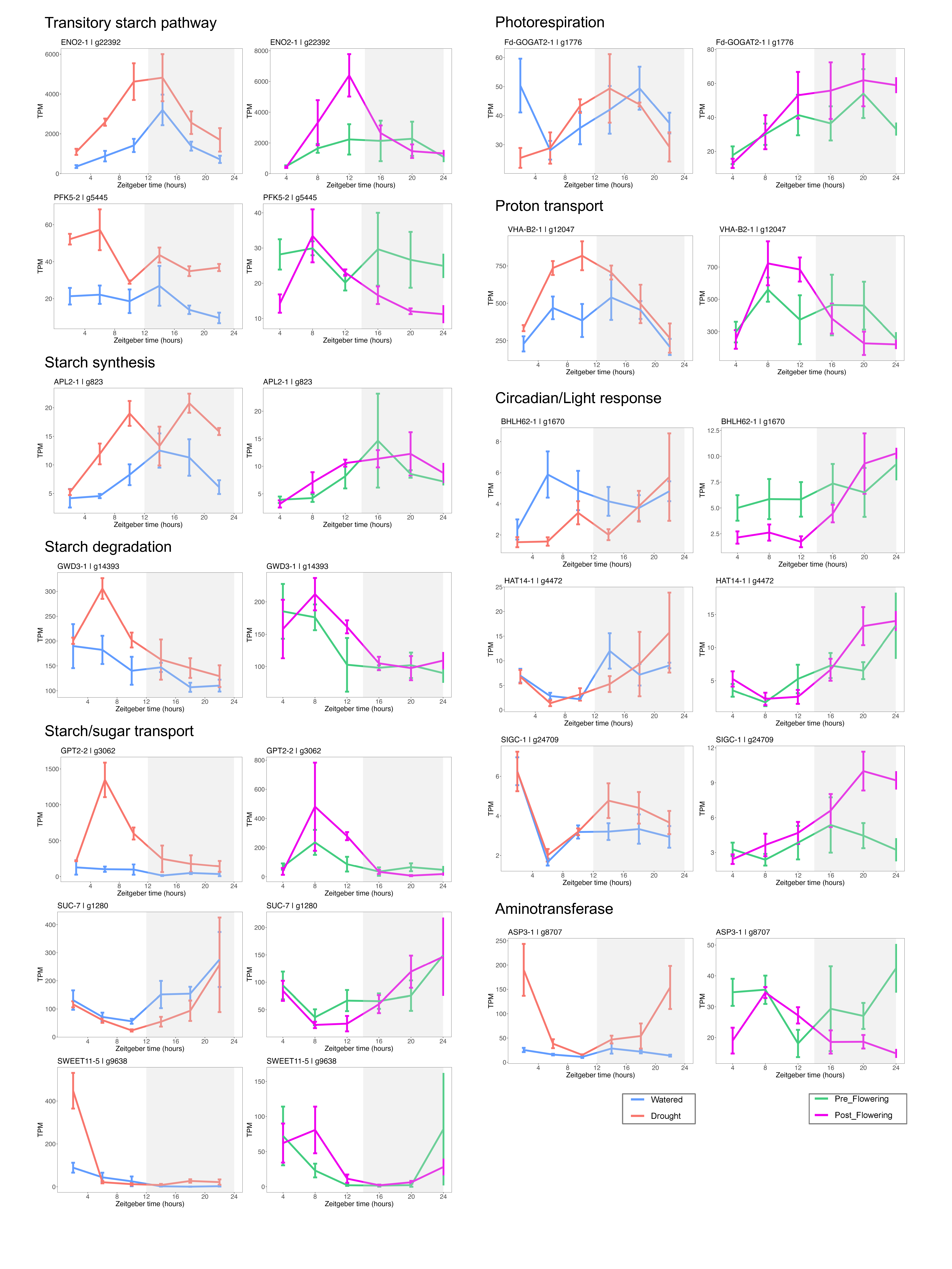

**Figure S17.** **Expression profiles of photosynthetic pathway genes with significant diurnal expression in both drought treatment and post-flowering stage, but not in watered treatment or pre-flowering stage**. Only plots of genes not previously shown are presented here. Normalized abundance of these genes are plotted across time (in TPM, transcripts per million), and grouped by photosynthetic pathway identity. Three or four biological replicates at each timepoint; error bars indicate interquartile range.

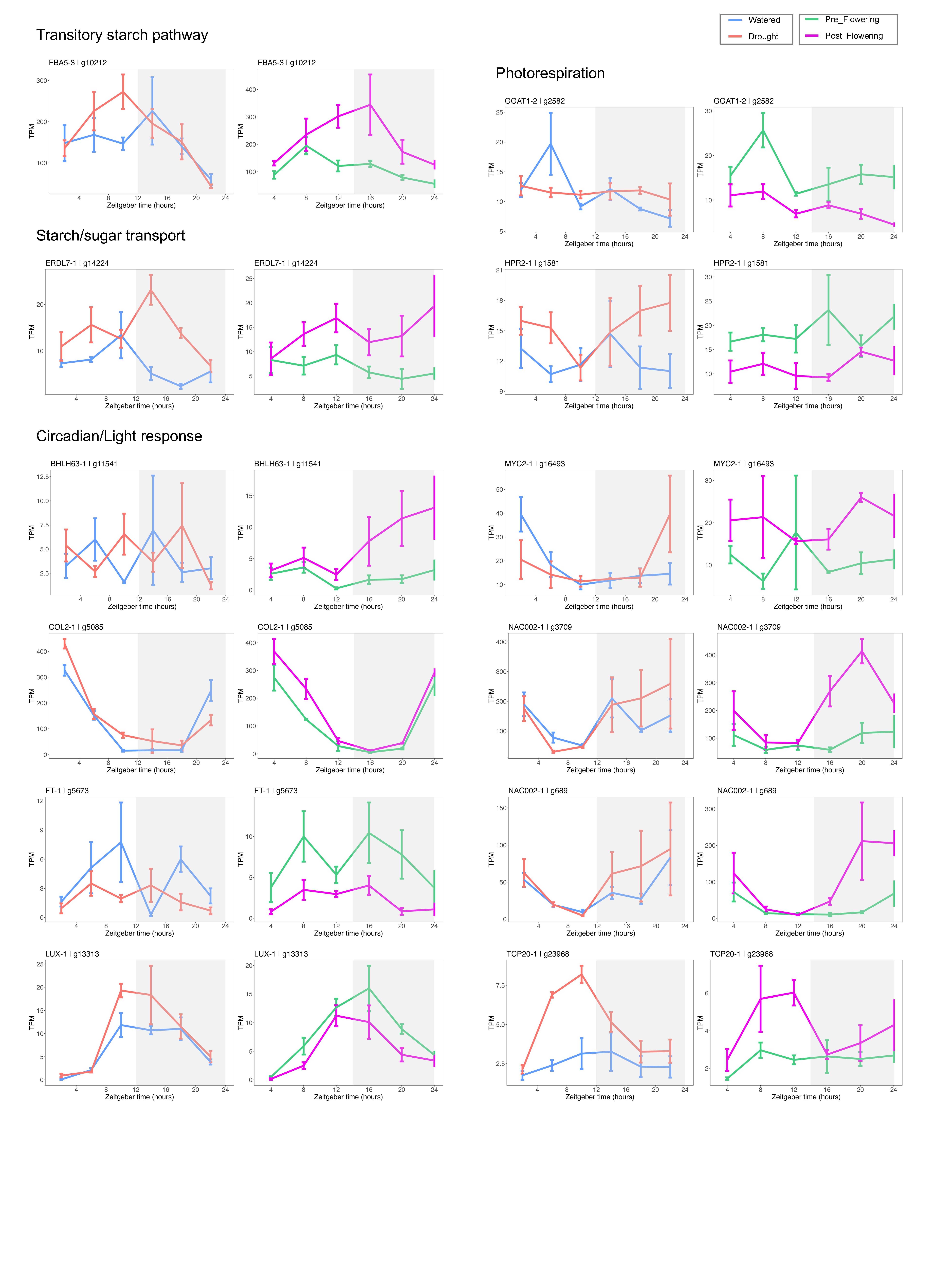

**Figure S18.** **Expression profiles of photosynthetic pathway genes with significant differential expression between Post-Flowering and Pre-Flowering stages, but not between Drought and Watered treatments.** Only plots not previously shown are presented here. Normalized abundance of these genes are plotted across time (in TPM, transcripts per million), and grouped by pathway identity. Three or four biological replicates at each timepoint; error bars indicate interquartile range.

**Table S1. *Cistanthe longiscapa* leaf samples collected across six time points for RNASeq and corresponding titratable acidity (ΔH^+^) values.** TRUE, sample was successfully sequenced; N/A, sample did not meet concentration and/or RIN score requirements (RIN ≥ 8) for library preparation; OR, sample concentration was out of range post-PCR prior to sequencing; SeqFail, sample was sequenced but failed to map successfully to reference genome.

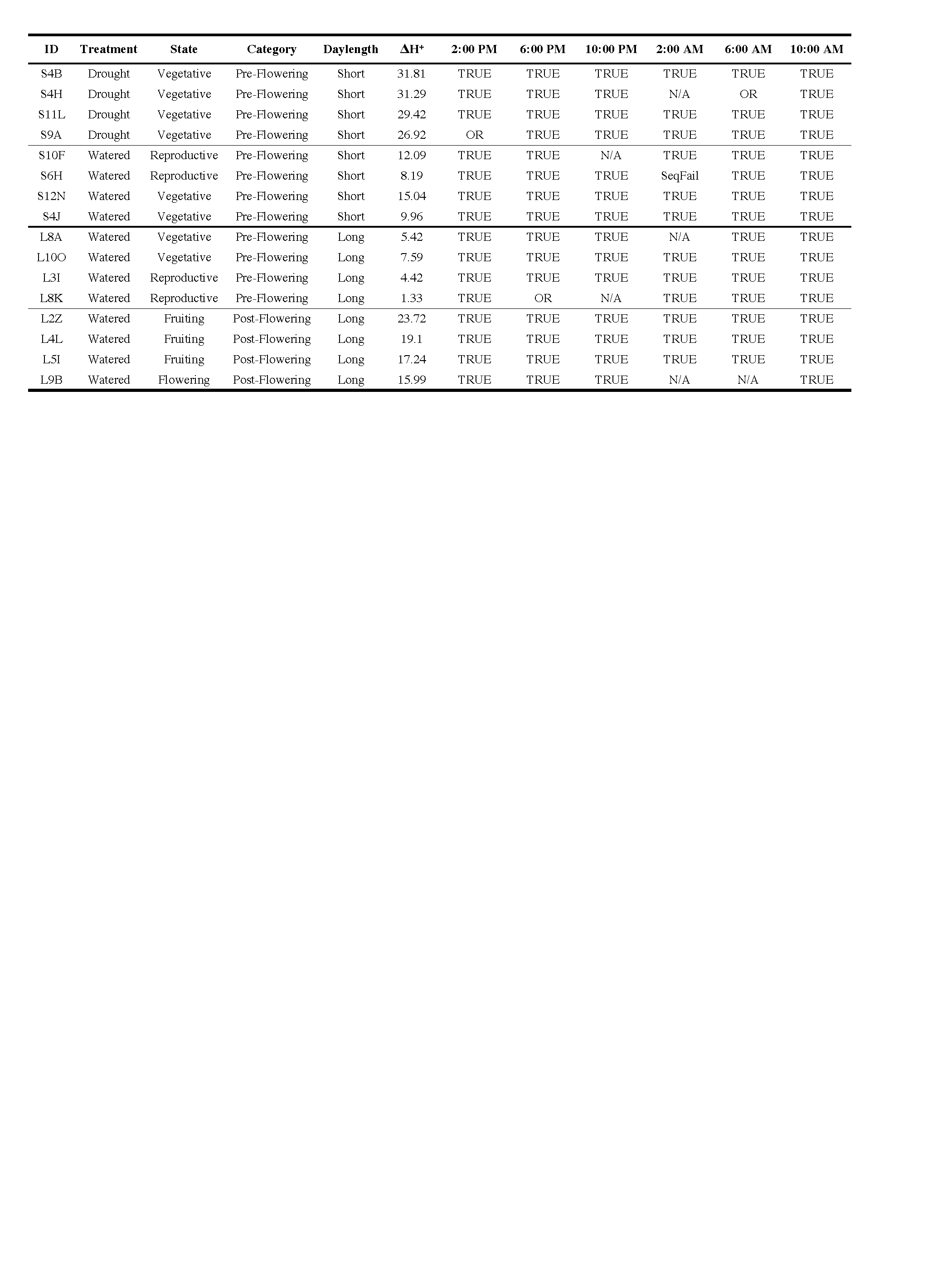

**Table S2. Detailed breakdown of repetitive elements of the *Cistanthe longiscapa* genome.** LINE, Long Interspersed Nuclear Element; LTR, Long Terminal Repeat; SINE, Short Interspersed Nuclear Element; TIR, Terminal Inverted Repeat.

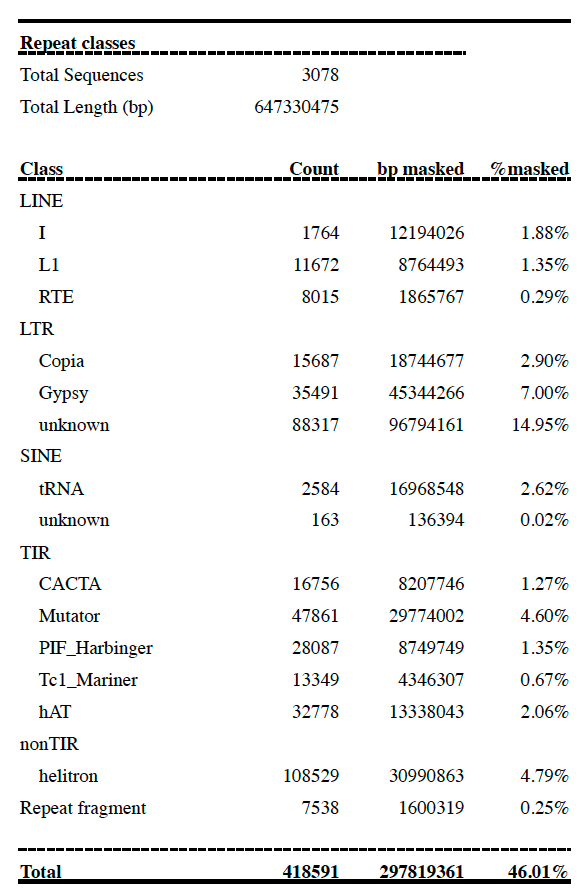
